# Principles of 3D Compartmentalization of the Human Genome

**DOI:** 10.1101/2020.11.22.393496

**Authors:** Michael H. Nichols, Victor G. Corces

**Affiliations:** Department of Human Genetics, Emory University School of Medicine, 615 Michael St, Atlanta, GA 30322, USA

**Author notes:** Corresponding author: Victor G. Corces, Department of Human Genetics, Emory University School of Medicine, 615 Michael Street, Atlanta, GA 30322.

**Keywords:** 3D organization, Chromatin, Transcription, Enhancer, Nucleus, CTCF, Cohesin

## Abstract

Chromatin is organized in the nucleus by CTCF loops and compartmental domains, the latter of which contain sequences bound by proteins capable of mediating interactions among themselves. While compartmental domains are one of the most prominent features of genome 3D organization at the chromosome scale, we lack a nuanced understanding of the different types of compartmental domains present in chromosomes and a mechanistic knowledge of the forces responsible for their formation. In this study, we compared different cell types to identify distinct paradigms of compartmental domain formation in human tissues. We identified and quantified compartmental forces correlated with histone modifications characteristic of transcriptional activity as well as previously underappreciated roles for compartmental domains correlated with the presence of H3K9me3, H3K27me3, or none of these histone modifications. We present a simple computer simulation model capable of predicting compartmental organization based on the biochemical characteristics of independent chromatin features. Using this computational model, we show that the underlying forces responsible for compartmental domain formation in human cells are conserved and that the diverse compartmentalization patterns seen across cells are due to differences in chromatin features. We extend these findings to *Drosophila* to suggest that the same fundamental forces are at work beyond humans. These results offer mechanistic insights into the fundamental forces driving the 3D organization of the genome.

## Introduction

The highly organized nature of the eukaryotic nucleus has been evident since experiments using immunofluorescence microscopy to determine the subnuclear distribution of various proteins and histone modifications showed the existence of several types of nuclear bodies (Matera et al., 2009). These nuclear locations, where proteins with related functional properties accumulate, have been described more recently as biomolecular condensates created as a consequence of liquid-liquid phase separation due to the presence of high concentrations of multivalent proteins bound to DNA and RNA, dividing the nucleoplasm into functionally distinct compartments (Banani et al., 2017). Some of these nuclear bodies appear to be involved in RNA-processing or sequestration, but others, such as the nucleolus, contain chromatin. These bodies represent distinct nuclear environments that regulate exposure of the DNA to various proteins of the nucleoplasm and are therefore essential to controlling the activity of genes. For example, active genes can be present in hubs termed transcription factories where expressed genes aggregate together with the transcriptional machinery (Mitchell and Fraser, 2008). Features of chromatin that are associated with transcriptional silencing also cluster with each other. Polycomb bodies form via the agglomeration of PRC1 and PRC2 protein complexes that epigenetically silence genes, in part by the trimethylation of H3K27 (Pirrotta and Li, 2012). Additionally, transcriptionally silenced pericentric heterochromatin colocalizes within the nucleus to form chromocenters in some cells, strongly enriched for HP1a and H3K9me3 (Wang et al., 2019). Several studies have now shown the ability of chromatin components to drive liquid-liquid phase separation in vitro and in vivo. H3K9me3 and HP1 together produce compartmentalization of heterochromatin, and this is also the case for the intrinsically-disordered regions found in PRC1, RNA Polymerase II and many transcription factors (Boijja et al., 2018; Ladouceur et al., 2020; Plys et al., 2019; Wang et al., 2019).

With the advent of Hi-C it has become possible to query the organization of the entire genome simultaneously at the sequence level (Lieberman-Aiden et al., 2009). Hi-C identifies all interactions in the genome after fixation with formaldehyde. The resulting contact frequency maps prominently display self-associating domains formed by short-range interactions among contiguous segments of the genome and can be visualized as “triangles” present at the diagonal of Hi-C heatmaps. Classically, these contact domains are called compartments and Topologically Associating Domains (TADs). The difference between these two types of domains is not functional but rather refers to the computational tools used to define them. Compartments are defined by Principal Component Analysis (PCA), normally using Hi-C data binned at 0.5-1.0 Mb resolution and, as a consequence, compartments are normally considered to be larger than 1 Mb in size. Compartments can contain sequences in an active (A) or silenced (B) transcriptional state and they interact with other compartments in the same state to give the plaid pattern observed in Hi-C heatmap. The term “compartment” is used to refer to both the self-interacting contact domains present at the diagonal as well as the ensemble of all the inter-domain interactions among all the domains in the same transcriptional state. To avoid confusion, we will use the term “compartmental domains” to refer to self-interacting contact domains present at the diagonal of Hi-C heatmaps and “compartment” to refer to all interactions among compartmental domains in the same transcriptional state. Different from compartmental domains, TADs are defined using algorithms that detect switches in the directionality of interactions. Analysis of Hi-C data at 1 kb resolution indicates that TADs actually correspond to two different types of domains--CTCF loops and compartmental domains (Rao et al., 2017; Rowley et al., 2017). CTCF loops are formed by the interruption of cohesin extrusion due to the presence of convergent CTCF-bound sites. CTCF loops can be visualized in Hi-C heatmaps by strong punctate signals at the summit of the domain, whereas compartmental domains lack this signal. CTCF loops disappear from Hi-C heatmaps obtained in cells depleted of CTCF or RAD21, whereas compartmental domains remain (Nora et al., 2017; Rao et al., 2017). Furthermore, compartmental domains are present in regions of the genome containing sequences in the same active or inactive transcriptional state and can be identified by PCA using 5-10 kb bin sizes. Neighboring regions in separate compartmental domains interact less frequently and represent a compartmental switch or border. In this way, the compartmentalization of the genome creates both local compartmental domains and distant compartmental interactions. Domains referred to as TADs in the literature are either compartmental domains, CTCF loops, or a combination of the two (Rowley and Corces, 2018).

As described above, compartmental domains can be captured by PCA of the Pearson correlation maps of each chromosome. The first principal component (PC1) or eigenvector captures the dimension with the highest variance. This vector has been divided into two sets of values because the corresponding sequences in the genome correlate well with transcriptionally active and inactive regions, and so are called A and B, respectively (Lieberman-Aiden et al., 2009). This categorization performs well across mammalian cell types and thus the compartmentalization of the genome is generally thought of as binary. However, epigenetic information suggests that the transcriptional state of the genome is more complex, and that the two-state classification is an oversimplification. While PCA remains the standard and most common method for calling compartmental domains, some analyses have more closely examined compartmentalization using more sophisticated techniques. The use of a Hidden Markov Model to cluster interchromosomal interactions resolved the active A and inactive B compartments into 2 active and 4 inactive compartments subtypes in GM12878 cells. These subcompartments differ according to various chromatin features, including post-translational histone modifications, replication timing, and measures of nucleolar and lamin association (Rao et al., 2014). However, the subcompartments within A and B do not differ substantially in which chromatin features are enriched. Other studies have also successfully used polymer simulations to reproduce the compartmental organization of the genome using chromatin-defined states (Annunziatella et al., 2018; Buckle et al., 2018; Chiang et al., 2019; Cook and Marenduzzo, 2018; Falk et al., 2019; Nuebler et al., 2018; Qi and Zhang, 2019). PCA-derived compartment calls in diverse cell lines and tissues invariably find A/B compartmentalization patterns, but the epigenetic features enriched in those A/B patterns can differ between cell-types. Notably, several studies have found the heterochromatin associated histone modification H3K9me3 strongly enriched in B compartments (Dixon et al., 2015; Falk et al., 2019). However, this modification is only found enriched in the single B4subcompartment in GM12878 cells, which is predominantly found only on chromosome 19 (Rao et al., 2014). Therefore, the binary A/B compartmentalization of the genome is far simpler than what would be predicted from microscopy analyses of nuclei where a large variety of biomolecular condensates composed of different epigenetic features have been observed.

Here we examine two cell types, GM12878 and HCT-116, with divergent compartmental definitions in an attempt to better understand the different patterns of compartmental domains seen between different human cells, and investigate the potential role of the forces responsible for this aspect of genome organization, with the goal of reconciling observations derived from Hi-C analyses and microscopy-based studies. The results suggest a consistent model of genome organization and offers insights into the mechanistic underpinnings of 3D genome compartmentalization.

## Results

### Dynamic compartmentalization across human cells

To better understand the mechanisms underlying the formation of compartmental domains and their compartmental interactions, we compared high-resolution Hi-C datasets containing 2.5-5 billions of contacts from two cell types – the lymphoblastoid GM12878 cell line and colorectal carcinoma HCT-116 cells. We took the Pearson correlation of the distance-normalized interaction maps in order to display the correlation of interactions of each bin with each other bin. This method can be used to visualize compartmental domains and their interactions because genomic regions in the same compartment will have highly correlated interaction frequencies and will have a high score in the correlation map. Figures 1A and 1B show the Pearson correlation maps for chromosome 4 of GM12878 and HCT-116, respectively. This chromosome shows very different organizations between the two cell lines. Both possess clear compartmental domains along the diagonal of the map and compartmental interactions as seen by the plaid pattern away from the diagonal. However, the locations and strength of these compartmental domains and the distal compartmental interactions in chromosome 4 from these two cell lines are very different. We sought to explore whether differences in the epigenetic profiles of the chromosomes of these two cell types could explain their distinct compartmentalization patterns. We compared the distribution of H3K27ac, which is correlated with transcriptional activity, H3K27me3, which is correlated with transcriptional silencing, and H3K9me3, which is also correlated with transcriptionally inactive sequences, to the Pearson correlation maps in these cells. We also performed PCA at 100 kb resolution and show the first principal component (PC1) in both cell lines (Figures 1A,B). Remarkably, GM12878 and HCT-116 show very different distributions of H3K9me3. In HCT-116 cells, large H3K9me3-rich domains correlate with prominent compartmental domains that are strongly correlated with each other in their Hi-C interaction frequencies. GM12878, on the other hand, lacks these large H3K9me3-rich domains and correspondingly lacks the prominent compartmental domains associated with them.

**Figure 1.**
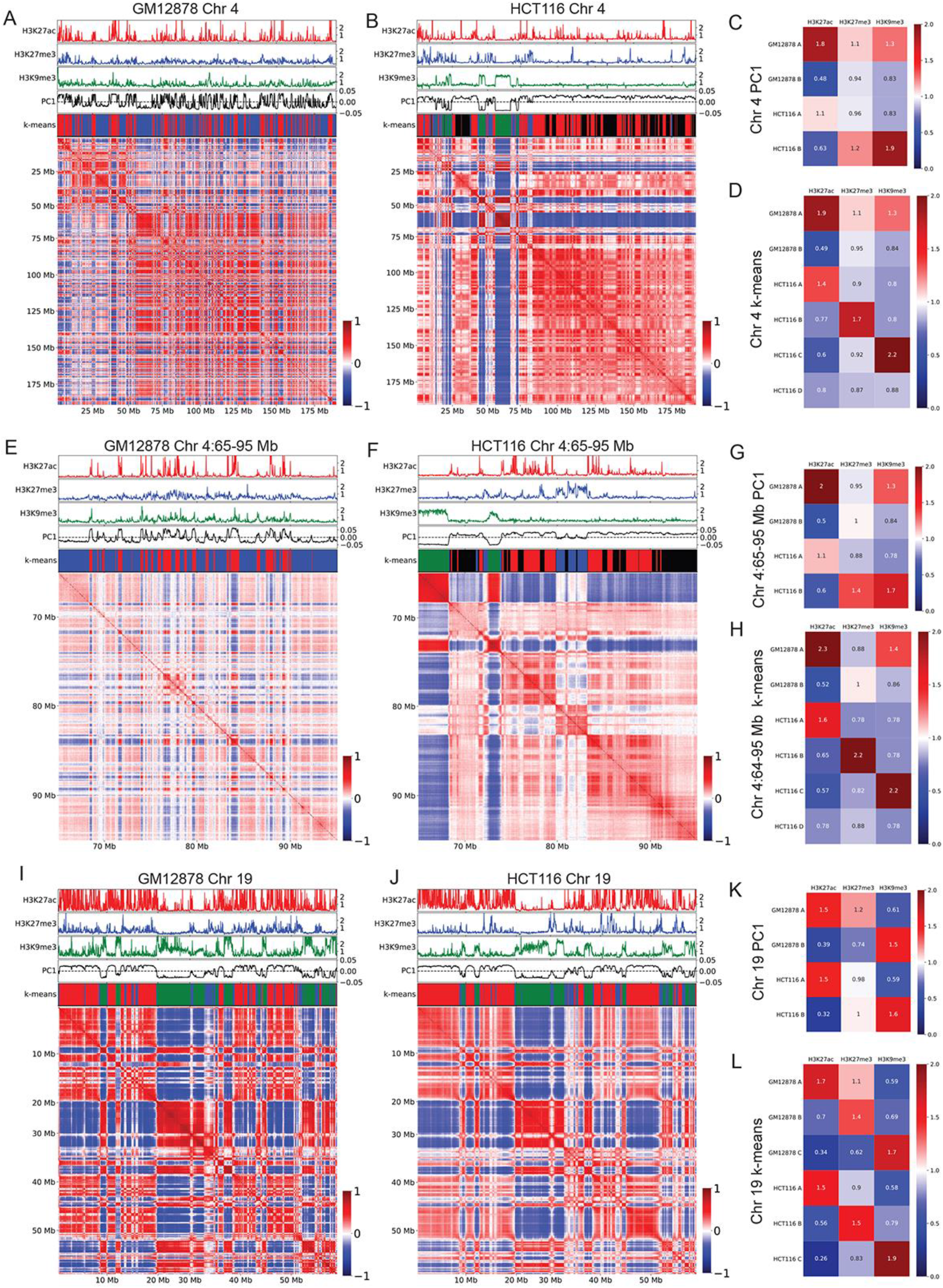
GM12878 and HCT-116 cells show different compartmental patterns. Pearson correlations of distance normalized Hi-C interaction frequency maps corresponding to various chromosome regions in GM12878 and HCT-116 cells. On top of each Hi-C map from top to bottom: fold-change over control shown for H3K27ac (red), H3K27me3 (blue), H3K9me3 (green); PC1 component from PCA (black), and k-means cluster classifications corresponding to compartments A (red), B (blue), C (green), D (black). A) Chromosome 4 from GM12878 cells. B) Chromosome 4 from HCT-116 cells. C) Fold-enrichment of each histone modification within each compartment defined by PCA in chromosome 4 from GM12878 and HCT-116 cells. D) Fold-enrichment of each histone modification within each compartment defined by k-means clustering on chromosome 4 from GM12878 and HCT-116 cells. E) Region spanning 65-95 Mb of Chromosome 4 from GM12878 cells. F) Region spanning 65-95 Mb of Chromosome 4 from HCT-116 cells. G) Fold-enrichment of each histone modification within each compartment defined by PCA on the region spanning 65-95 Mb of chromosome 4 from GM12878 and HCT-116 cells. H) Fold-enrichment of each histone modification within each compartment defined by k-means clustering on the region spanning 65-95 Mb of chromosome 4 from GM12878 and HCT-116 cells. I) Chromosome 19 from GM12878 cells. J) Chromosome 19 from HCT-116 cells. K) Fold-enrichment of each histone modification within each compartment defined by PCA on chromosome 19 from GM12878 and HCT-116 cells. L) Fold-enrichment of each histone modification within each compartment defined by k-means clustering on chromosome 19 from GM12878 and HCT-116 cells.

To quantify these observations, we used PC1 to call A and B compartmental domains in GM12878 and HCT-116 cells and measured the relative enrichments of chromatin features on their chromosomes (Figure 1C). In GM12878 cells, A/B compartmentalization strongly follows transcriptional activity/inactivity, with histone modifications associated with active transcription such as H3K27ac, H3K4me3, and H3K36me3 all enriched in A compartmental domains and depleted in the B compartment of chromosome 4, while modifications associated with silenced chromatin such as H3K27me3 show an inverse pattern (Figure 1C and Supplemental Figure 1A). In HCT-116 cells, however, A compartmental domains in chromosome 4 are not strongly enriched for histone modifications associated with transcriptional activity, including Gro-seq, which is a direct measure of transcription. Instead, binary compartmental delineation using PC1 divides the chromosome into a transcriptionally inactive H3K9me3-rich portion and the remainder, which consists of both transcriptionally active as well as inactive regions enriched in H3K27me3. In contrast, H3K9me3 in chromosome 4 of GM12878 is enriched in A compartmental domains and presents very differently on the chromosome as sporadic peaks rather than contiguously enriched domains (Figure 1C and Supplemental Figure 1A). Given that B compartmental domains in chromosome 4 of HCT-116 cells are depleted of transcriptionally active sequences, we asked why the corresponding A compartmental domains are not strongly enriched for transcribed sequences. A simple explanation for this phenomenon is that transcriptionally inactive regions of HCT-116 not containing H3K9me3 correlate more closely in their interaction frequencies with transcriptionally active regions. Thus, the A compartment in these cells as defined by PCA is composed of a conglomeration of all H3K9me3-poor chromatin, both transcriptionally active and inactive, leading to only a mild enrichment for active histone modifications. These observations differ from the general assumption that A compartmental domains defined by PCA correspond exclusively to actively transcribed sequences and may explain the observed lack of correlation between changes in transcription and compartmental switches.

To examine more closely the mechanisms underlying the formation of compartmental domains we focused on a 65-95 Mb region on chromosome 4 and used a resolution of 25 kb to call compartments using PCA (Figures 1E,F). In HCT-116 cells, this region contains instances of all 4 clusters found in chromosome 4. Compartmental domains present in the A compartment defined by PCA in GM12878 cells are highly enriched in H3K27ac with respect to those in the B compartment, whereas H3K27me3 and H3K9me3 are similarly enriched in both A and B compartments (Figures 1E,G). However, in HCT-116 cells there is a clear enrichment of both H3K9me3 and H3K27me3 in the B compartment corresponding to compartmental domains and interactions absent in GM12878 cells (Figures 1F,G). These findings, showing differential enrichment of active and repressive histone modifications in the A and B compartments in different cell lines, are surprising, since it is generally assumed that the A compartment contains transcriptionally active genes and the B compartment is enriched in silenced sequences independent of the cell type. This suggests that the canonical binary classification of A and B compartments is insufficient to represent the properties and mechanisms by which compartmental domains form in these cells.

### Conserved principles underlie dynamic compartmentalization

To further explore the complex compartmentalization logic observed in GM12878 and HTC-116 cells, which appears to follow different rules in the two cell lines and cannot be explained by a simple binary division of PC1, we employed the unsupervised k-means clustering algorithm to identify compartmental clusters in both cell types. The primarily binary A/B organization of chromosome 4 in GM12878 cells obtained by PCA can be reproduced by two clusters obtained via unsupervised k-means clustering (Figures 1A,E). However, four clusters are required to produce a meaningful classification of chromosome 4 that correlates with the Hi-C heatmap in HCT-116 cells (Figures 1B,F). As expected, one of these four clusters corresponds directly to H3K9me3-rich regions whereas a second one correlates strongly with H3K27ac. Surprisingly, the two other clusters are both transcriptionally inactive with distinct chromatin features, one highly enriched for H3K27me3, and the last lacking all three histone modifications (Figures 1D and 1H). Only H3K9me2 is enriched in this compartment (Supplemental Figure 1A). Therefore, chromosome 4 in HCT-116 appears to have four distinct compartmental domains. Interactions among each type give rise to the complex plaid pattern in the Hi-C heatmap, forming compartment A (transcriptionally active), B (H3K27me3-rich), C (H3K9me3-rich), and D (enriched in H3K9me2 but depleted of standard active and silencing histone modifications). This 4-compartment nomenclature corresponding to enrichment in each of these different histone modifications will be used in the rest of the manuscript. We note that the enrichments of epigenetic features in chromosome 4 A and D clusters of HCT-116 cells correspond well to the A and B compartments of chromosome 4 in GM12878 cells, with the exception of H3K9me3 enrichment in the A compartment of GM12878 (Figures 1C,D).

We then sought to understand why H3K9me3 is enriched in different compartmental domains in HCT-116 versus GM12878 cells. The distribution of H3K9me3 in these two cell types is very different, with GM12878 chromosomes typically having narrow peaks of signal whereas HCT-116 chromatin tends to have large consistently enriched plateaus. At least two possible hypotheses could explain these different distribution patterns. One possibility is that H3K9me3 regions compartmentalize differently in the two cell types due to different nuclear environments determined by cell identity and physiology. A second explanation is that H3K9me3 regions compartmentalize differently due to distinct distributions of this histone modification as a consequence of transcriptional differences between the two cell types. Analysis of chromosome 19 offers an opportunity to distinguish between these two possibilities, since this chromosome contains large domains of H3K9me3 in both GM12878 and HCT-116 cells. Chromosome 19 of GM12878 cells shows a strong correlation between the presence of H3K9me3 and the formation of strong compartmental domains (Figure 1I), as was seen on chromosome 4 of HCT-116 cells, and these H3K9me3 compartmental domains are similar in chromosome 19 of both cell types (Figure 1J). The similarity between H3K9me3 domains in chromosome 19 of GM12878 and HCT-116 cells suggests that the nuclear environment is not responsible for the differences observed in other chromosomes. Strikingly, the histone modification profiles and the A/B compartments defined by PC1 also closely match between the two cell lines (Figure 1K). k-means clustering was then performed in both cell-types with 3 clusters, since chromosome 19 lacks large regions devoid of any signals that would fall into the D cluster. The resulting cluster calls closely match each other (Figure 1L). The high correspondence in k-means cluster definitions is mirrored by the relative signal enrichments in each cluster. In contrast to chromosome 4, the compartments called by PCA and by k-means showed similar enrichments for various histone modifications in chromosome 19 (Figure 1L and Supplemental Figure 1C). The alignment of chromosome 19 in histone modification profiles and compartmental organization in HCT-116 and GM12878 fits a model where these histone modifications, or a chromatin feature correlated to them, drive chromosomal compartmentalization, and suggests that the underlying forces driving compartmentalization are consistent between these cell types. We suggest that the diverging compartments seen in chromosome 4 are the result of distinct chromatin profiles and that the resulting conflicting histone modification enrichments for the A and B compartments of chromosome 4 between these cell lines do not reflect differences in the underlying chemistry driving compartmentalization. The enrichment of H3K9me3 in the active compartment of GM12878 chromosome 4 can be explained as the inability of small narrow peaks of H3K9me3 to drive compartmentalization against entropic mixing. Their proximity to other active compartmentalizing features may further inhibit their self-segregation. Alternatively, since these short regions containing H3K9me3 are much smaller than the 25 kb bins used to perform the clustering analyses, it is possible that compartmental calls performed at a higher resolution might resolve H3K9me3-associated compartmentalization in these small regions. This possibility highlights the potential biases introduced by the resolution of the data used in the analyses when interpreting Hi-C information and the need for improved conformation data.

We then considered whether either of these cell lines perhaps exemplify an outlier unrepresentative of normal human tissues and thus we investigated H3K9me3 distribution in numerous immortalized and primary human cells. We found that neither the H3K9me3 pattern observed in GM12878 and HCT-116 cells is unrepresented in other cell types or in primary tissues but rather exemplifies two ends of a continuum of H3K9me3 present across human cells (Supplemental Figure 1D, E). These findings reveal that H3K9me3 domains are highly dynamic and give rise to significant changes in compartmentalization patterns across development.

### Compartmental domains correlate directly with chromatin features

With the understanding that binary models are insufficient to represent human compartmentalization and that the forces responsible for the formation of compartmental domains appear to correlate closely with chromatin features, particularly transcriptional activity, H3K27me3, and H3K9me3, we sought to more precisely visualize and quantify the forces driving compartmentalization. To this end, we used chromosome 14 from HCT-116 cells as an example and sorted the Pearson correlation map by various features (Figure 2A). These assemblies are produced by reordering the rows and columns of the correlation matrix so that, instead of being placed in their natural order, bins are arranged by increasing signal of the chosen feature. This allows us to compare the ability of a particular sequence to form a specific type of compartmental domain with its epigenetic features. This approach also provides a means of visualizing the frequency of interactions within and among compartmental domains or “compartmentalization strength”. Sorting by PC1 values we observe a clear segregation of chromosome 14 into three types of compartmental domains A, B, and C (Figure 2B). One type of compartmental domain-named compartment C above-corresponds to negative values of PC1 and contains sequences enriched in H3K9me3 but lacking H3K27me3 and H3K27ac. A second type of compartmental domain-compartment B-corresponds to intermediate positive PC1 values and contains sequences enriched in H3K27me3 but lacking H3K9me3 and H3K27ac. Finally, a third A compartment contains sequences with high positive PC1 values, and it is enriched in sequences with high levels of H3K27ac but lacking the other two silencing histone modifications. We note that A and C compartmental domains are highly self-correlated but strongly anticorrelated with each other in interaction frequency while B compartmental domains self-correlate and have intermediate levels of correlation with both A and C domains (Figure 2B). B compartmental domains have higher correlation with A than with C domains, indicating they interact more similarly to A and would be grouped with A by a binary classification system (Figure 2B). The existence of these three compartments is supported by results from k-means clustering (see green/blue/red bar in Figure 2B and other panels). Surprisingly, this indicates that PC1, while typically used to delineate binary compartments, could be used to call more compartmental domain types with different thresholds. Chromosome 14, like chromosome 19, largely lacks regions that would fall into the D compartment. Sorting the chromosome by H3K9me3 instead of PC1 values reproduces the compartmentalization of the H3K9me3-rich C compartment as well as the first principal component itself, however, it was unable to distinguish between the A and B compartments (Figure 2C). Sorting by H3K27ac as a marker of transcriptional activity also results in compartmental segregation, although this is less precise (Figure 2D). This is likely a function of H3K27ac signal lacking the consistency and continuity of H3K9me3 domains within transcriptionally active regions. Sorting by H3K27me3 organized only the regions most enriched for H3K27me3 and was unable to organize the rest of the chromosome (Figure 2E). Together these results show that covalent histone modifications strongly correlate with and are predictive of compartmental organization.

**Figure 2.**
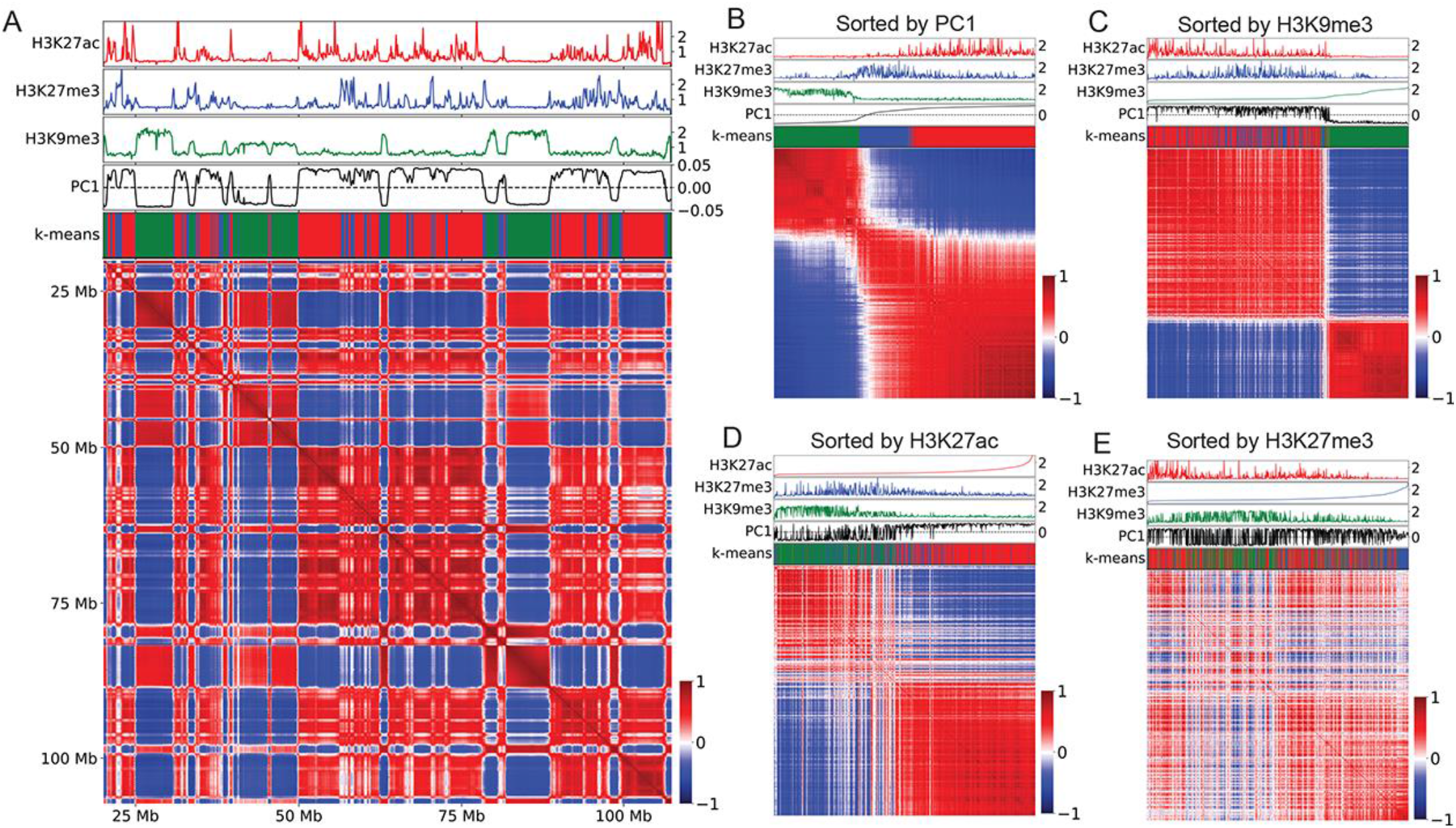
Results from sorting sequences from chromosome 14 of HCT-116 cells based on the magnitude of PC1 and the levels of various histone modifications. Pearson correlations of distance normalized Hi-C interaction frequency map of chromosome 14 from HCT-116 cells. Shown above each Hi-C map from top to bottom are the following: fold-change over control for H3K27ac (red), H3K27me3 (blue), H3K9me3 (green); PC1 (black), and k-means cluster classifications A (red), B (blue), C (green). A) Interactions among sequences from chromosome 14 arranged in its natural order. B) Interactions among sequences from chromosome 14 sorted according to PC1 values from lowest to highest. C) Interactions among sequences from chromosome 14 sorted according to H3K9me3 levels from lowest to highest. D) Interactions among sequences from chromosome 14 sorted according to H3K27ac levels from lowest to highest. E) Interactions among sequences from chromosome 14 sorted according to H3K27me3 levels from lowest to highest.

An unusual feature of chromosome 14 is the existence of a trimodal distribution of H3K9me3. While most chromosomes in GM12878 do not show 100 kilobase regions strongly enriched for H3K9me3 (Supplemental Figure 2A and C), most chromosomes in HCT-116 cells possess a bimodal distribution of H3K9me3 signal leading to H3K9me3 rich and poor regions (Supplemental Figure 2B). In HCT-116, chromosomes 13 and 14 have distinct strong and weak H3K9me3 domains (Figure 2A, Supplemental Figure 2D). Comparing their correlations in the sorting of chromosome 14 by H3K9me3 shows that weak-H3K9me3 and strong-H3K9me3 regions correlate better with regions of similar strength (Figure 2C). This phenomenon, along with the varying correlation strength seen in the heatmaps obtained by sorting various features, suggests that compartmentalization is more accurately thought of as a quantitative rather than a categorical feature of the genome and that classification of chromatin into categories is a simplification that neglects to consider the effects of varying epigenetic signal strengths in the formation of compartmental domains. Sorting by a single signal does not result in perfect compartmentalization because multiple, independent forces drive the process of compartmental domain formation and long-range interactions among domains of the same type. The most significant of these forces may be those involved in interactions between sequences containing H3K9me3 and proteins and histone modifications associated with active transcription. These observations suggest a model of compartmentalization in which the genome is organized by multiple independent forces directly correlated with chromatin features that attract and repel each other.

### Independent contributions of chromatin features can reproduce compartmental organization

We next sought to test the hypothesis that formation of compartmental domains and establishment of long-range interactions among domains of the same type is driven by the independent contributions of chromatin features. To approach this question, we created a machine learning model to reproduce Hi-C interaction maps using epigenetic features. Results described above suggest that the presence of H3K27ac as an indicator of transcriptionally active regions, H3K27me3 and H3K9me3 as indicators of different types of silenced sequences, or the absence of these three modifications can account for all the compartmental domains we observed in chromosomes of human cells. To enable comparison across cell types and experiments, we first binned these three epigenetic signals into quantiles at 100 kb resolution. For each normalized signal, an algorithm then learned an attraction-repulsion relationship for each pair of quantiles using a Maximum Likelihood Estimation approach. This attraction-repulsion mapping effectively represents the average enrichment or depletion between all bins with the corresponding level of each chromatin feature. The estimated contact frequency in the simulated map is then derived by the simple addition of the estimated effect of each signal and multiplied by a distance-dependent constant representing the average interaction frequency at each genomic distance (Figure 3A). This relatively straightforward model, which is only capable of representing the independent attractive and repulsive forces of each chromatin feature, tests to what extent this framework is capable of recapitulating the 3D organization of the genome represented by the compartmental domains and their interactions.

**Figure 3.**
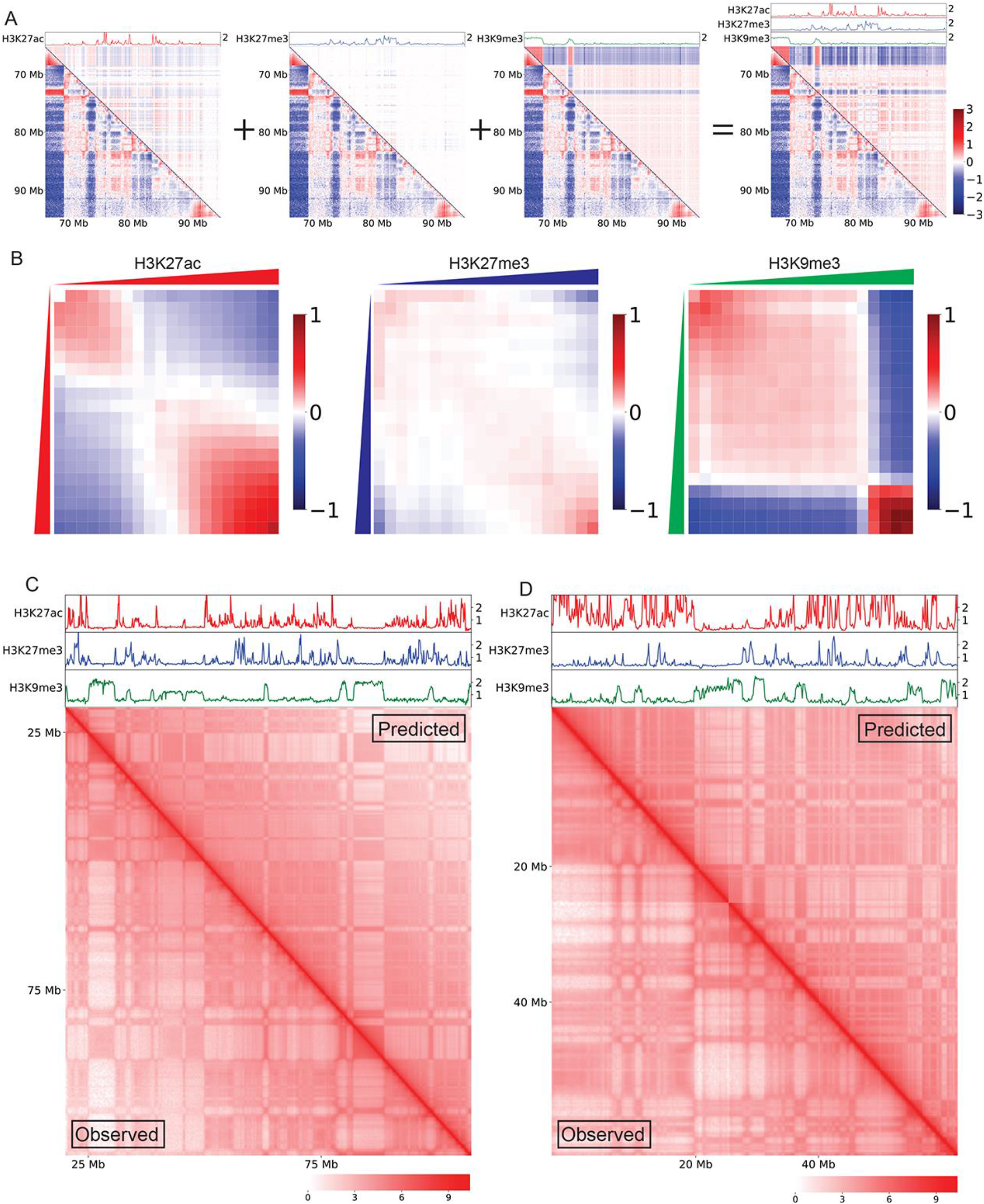
Histone modifications can predict compartmentalization using learned attraction-repulsion relationships. A) Log(observed/expected) of Hi-C interaction maps showing the 65-95 Mb region of chromosome 4 from HCT-116 cells. Bottom left triangles are observed Hi-C interaction maps while upper right triangles are simulations using only the components shown above as tracks. From left to right H3K27ac (red), H3K27me3 (blue), H3K9me3 (green), and all three combined. B) Average of attraction-repulsion relationship maps learned by Maximum Likelihood Estimation from every chromosome of HCT-116 cells. C) Comparison of HCT-116 chromosome 8 logged Hi-C interaction maps. The bottom left triangle corresponds to observed interactions and the upper right triangle represents simulated contacts. D) Comparison of HCT-116 chromosome 19 logged Hi-C interaction maps. The bottom left triangle corresponds to observed interactions and the upper right triangle represents simulated contacts.

The averaged attraction-repulsion relationships learned from every chromosome of HCT-116 cells are shown in Figure 3B. The learned relationship between the levels of a specific histone modification and the interaction frequency of the corresponding sequence is similar to that observed experimentally described above (Figures 2C-2E). Genomic regions containing high levels of a given histone modification show increased interactions with other regions high in the same modification. The model learns and predicts that pairs of regions in which one is high in such a signal and the other low will not be attracted and have reduced interaction frequency. The minimal model using three histone modifications, H3K27ac, H3K27me3, and H3K9me3, is able to recapitulate most aspects of 3D genome organization while remaining easy to interpret. All three signals show a degree of attraction between the highest quantile bins, as seen by the enrichment in the bottom right of the attraction-repulsion maps, as well as repulsion between the highest and lowest quantiles as seen by the depletion in the upper right and bottom left corners (Figure 3B). Importantly the strength and nature of these maps differ significantly, indicating the forces driving compartmentalization vary for each feature. H3K9me3 maps show strong attraction amongst the most enriched quarter of the genome, which strongly repels the rest of the genome equally. This shows the existence of a single critical threshold of H3K9me3 density and quantity, and reflects the generally bimodal distribution of this modification in HCT-116 cells. H3K27ac, on the other hand, shows the greatest attraction among the highest quantiles with a more gradual reduction in attraction with reduced signal. H3K27me3 primarily shows only attraction between the highest quantiles of signal and otherwise contributes little to the organization of the genome. When interpreting this information, it is important to consider that the distributions of these three histone modifications are inter-related, and that regions of the genome lacking one of the modifications may contain one of the other two.

Simulations were generated at 100 kb resolution using the average of the attraction-repulsion maps learned from every chromosome except the one being simulated. A comparison of observed and simulated maps reveals close agreement on the majority of large compartmental features (Figure 3C,D and Supplemental Figure 3A,B). We quantified the accuracy of the model using the Pearson correlations between the observed and simulated maps after dividing by the average distance. Due to the power-law decay of interaction frequency with respect to distance in Hi-C maps any simulation that accurately reproduces this decay will have a high correlation. As this would not represent the ability of the model to reproduce compartmental organization, we normalized for distance to eliminate the natural correlation driven by the accurate representation of the distance decay. Simulations of chromosomes using the average maps derived from all other chromosomes varied in correlation by chromosome, but generally performed well with correlations in the range 0.5-0.7 (Supplemental Figure 3C). Using this same methodology to compare biological replicates of a Hi-C experiment resulted in similar ranges of correlation scores across the chromosomes (Supplemental Figure 3D). The fact that Hi-C maps can be predicted by only modeling the attraction and repulsion of chromatin features against themselves suggests a direct role of these features, or a chromatin component correlated with these features, in compartmentalizing the nucleus.

### Conserved forces give rise to diverse genomic organizations

Given the accuracy of our model in reproducing the 3D organization of HCT-116 cells, we then applied the same model to GM12878 cells. These cells have different distribution of H3K9me3, and their 3D genome organization as seen in PCA enrichment analysis is also different (Figure 1C). We found that, with the exception of H3K9me3, the average of the attraction-repulsion maps learned across all chromosomes in GM12878 were strikingly similar to those learned in HCT-116 (Figure 4A). The incongruence of H3K9me3 attraction-repulsion maps between these cell types was expected as the distribution of H3K9me3 is very different between most of their chromosomes. We simulated the Hi-C interaction maps of GM12878 chromosomes using the average of the attraction-repulsion maps learned from all other chromosomes (Figure 4B, Supplemental Figure 4A,B). However, simulating chromosome 19 in GM12878, which is unique compared to other chromosomes in these cells due to its large H3K9me3 domains, was less successful when using the attraction-repulsion maps learned from the rest of the chromosomes, with the simulation failing to capture the role of H3K9me3 in compartmentalization (Figure 4C). Pearson correlations between the real and simulated maps were significantly lower in GM12878 cells than in HCT-116 indicating the simulation was less successful at reproducing the Hi-C maps in GM12878, most likely due to the absence of the strong organizing feature of H3K9me3 (Supplemental Figure 4C). Nevertheless, the similarity between the attraction-repulsion maps of these two cell types suggests that the fundamental forces underlying compartmentalization are largely conserved between these cells. If this is true, the attraction-repulsion maps learned in one cell type would be able to accurately model the organization of another cell type, showing that dynamic compartmentalization is largely a consequence of diverging distributions of chromatin features.

**Figure 4.**
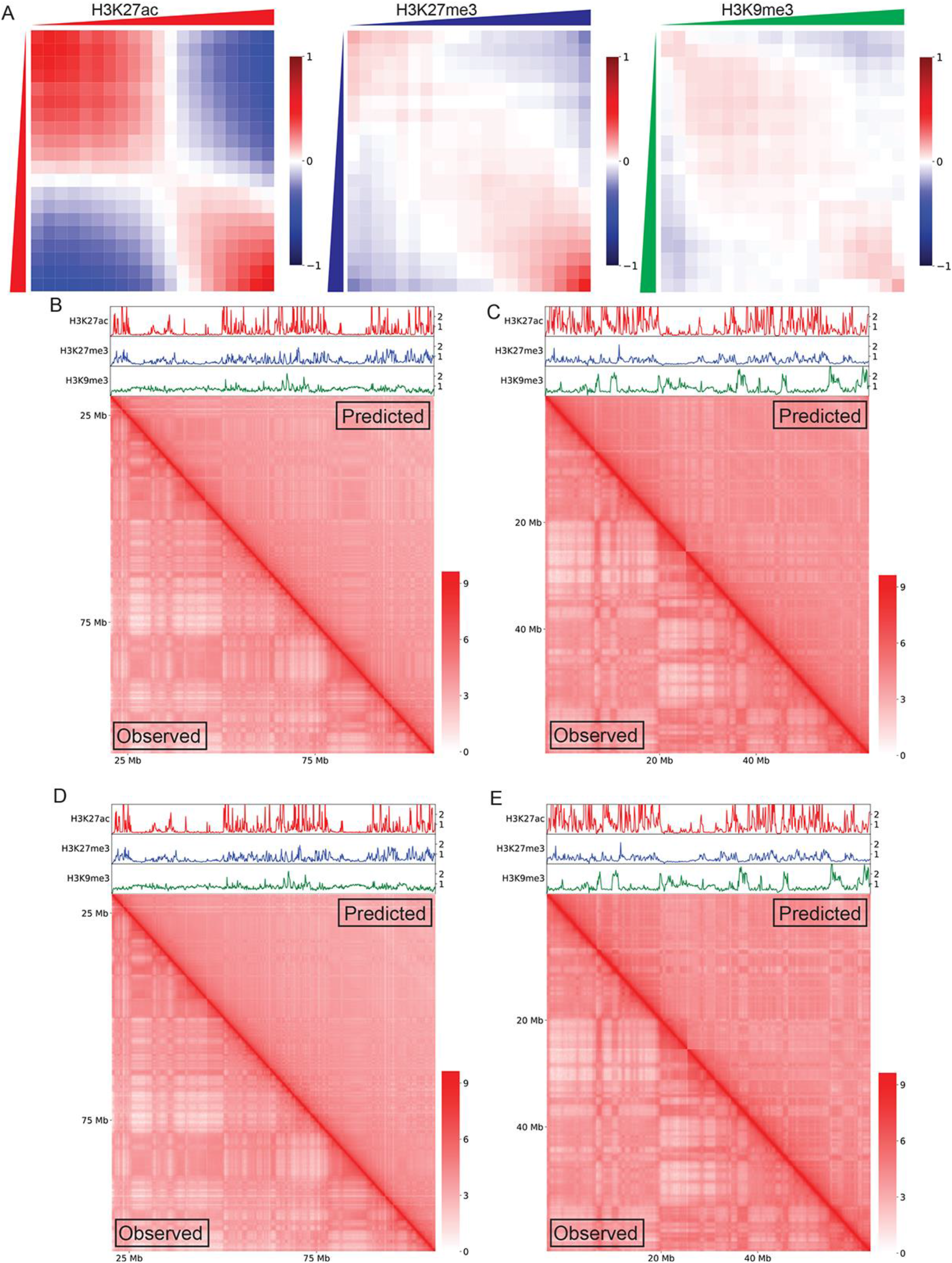
Attraction-repulsion relationships are consistent across cell types. A) Average of attraction-repulsion relationship maps learned by Maximum Likelihood Estimation from every chromosome of GM12878 cells. B) Comparison of GM12878 chromosome 8 logged Hi-C interaction maps. The bottom left triangle corresponds to observed interactions and the upper right triangle represents simulated contacts using attraction-repulsion maps learned from GM12878. C) Comparison of GM12878 chromosome 19 logged Hi-C interaction maps. The bottom left triangle corresponds to observed interactions and the upper right triangle represents simulated contacts using attraction-repulsion maps learned from GM12878. D) Comparison of GM12878 chromosome 8 logged Hi-C interaction maps. The bottom left triangle corresponds to observed interactions and the upper right triangle represents simulated contacts using attraction-repulsion maps learned from HCT-116. E) Comparison of GM12878 chromosome 19 logged Hi-C interaction maps. The bottom left triangle corresponds to observed interactions and the upper right triangle represents simulated contacts using attraction-repulsion maps learned from HCT-116.

To test this idea we used the attraction-repulsion maps learned from HCT-116 chromosomes to simulate the 3D organization of GM12878 chromatin, and found a high correspondence between the observed and simulated maps (Figure 4D, Supplemental Figure 4D,E,F). This is remarkable, given that the Hi-C interaction maps of most chromosomes between these cell types are very different. This ability for compartmental forces learned from one cell type to successfully predict compartmentalization in another indicates that the same underlying forces directly correlated with chromatin features are largely conserved between these two cell types. This is further support that the radical differences between the respective Hi-C maps of these cell lines are a consequence of different chromatin features, primarily the presence and absence of large H3K9me3-rich domains.

As the simulation of chromosome 19 of GM12878 cells using attraction-repulsion maps learned with the rest of the chromosomes is poor, we asked whether chromosome 19 could be better simulated by the attraction-repulsion relationships learned from HCT-116 than from GM12878. The histone modification profiles of chromosome 19 in GM12878 cells, particularly the presence of large H3K9me3 domains, more closely resemble the patterns seen in most chromosomes of HCT-116 cells (Figure 1G). Indeed, the simulation of chromosome 19 is significantly more accurate using HCT-116 attraction-repulsion maps (Figure 4E; compare with Figure 4C). In agreement, the Pearson correlations between the observed and simulated maps of chromosome 19 are higher when simulated with HCT-116 (Supplemental Figure 4F). These results again support the idea that the 3D organization of chromosomes is a consequence of the distribution patterns of one-dimensional epigenetic information.

### Transcriptional activity and H3K9me3 also compartmentalize the *Drosophila* genome

Given the ability of our model to reproduce the 3D organization of multiple human cell types from a single universal set of attraction-repulsion maps, we sought to determine the applicability of this model outside of humans. Our previous foundational work showed that transcriptional activity was predictive of compartmental organization in multiple representative Eukaryotic genomes (Rowley et al., 2017). We hypothesized that, just like in humans, the same forces responsible for compartmental domain formation and the establishment of distinct nuclear compartments via interactions among compartmental domains with the same epigenetic features may be at work in other organisms. Thus, we applied our model to explore whether the distribution of H3K27ac, H3K27me3, and H3K9me3 could reproduce the 3D organization of the *Drosophila* genome.

Hi-C maps from *Drosophila* Kc cells using datasets with approximately 1 billion reads were simulated using the attraction-repulsion model. Due to the higher read count and smaller genome size we were able to simulate the genome at 10 kb resolution. A small modification was made to the model to limit the simulation to less than 2 Mb, as *Drosophila* compartmental domains are smaller than those of mammals and long-range interactions among domains decay rapidly beyond this distance. While distinct, the learned attraction-repulsion maps resemble those learned in human cells (Figure 5A). The model learns attraction between features with similar histone modifications and repulsion between genomic regions with dissimilar ones.

**Figure 5.**
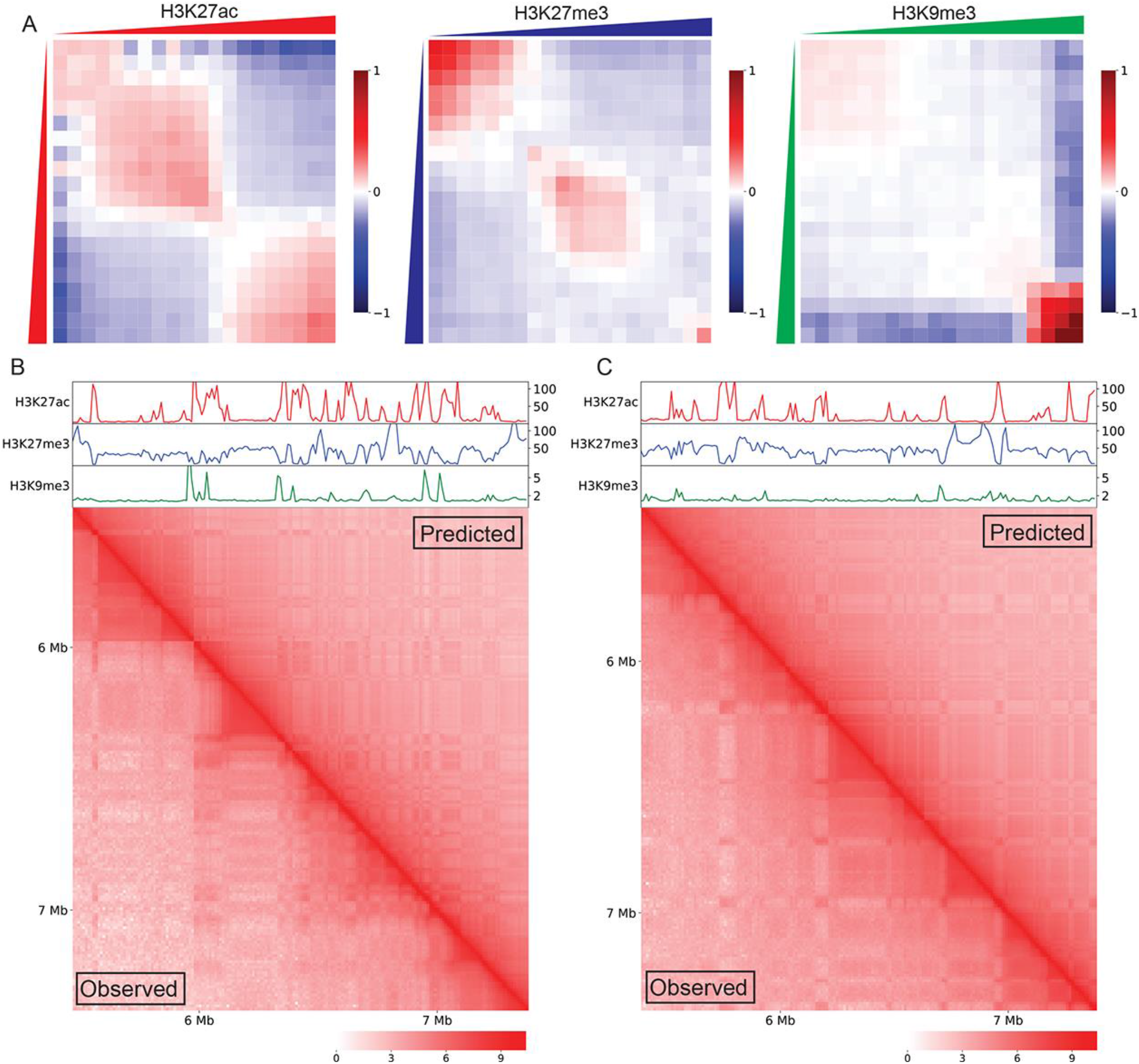
Attraction-repulsion relationships explain compartmentalization in *Drosophila*. A) Average of attraction-repulsion relationship maps learned by Maximum Likelihood Estimation from chromosomes 2 and 3 of Kc167 cells. B) Comparison of Kc167 chromosome 2 logged Hi-C interaction maps. The bottom left triangle corresponds to observed interactions and the upper right triangle represents simulated contacts using attraction-repulsion maps learned from chromosome 3 of Kc167 cells. C) Comparison of Kc167 chromosome 3 logged Hi-C interaction maps. The bottom left triangle corresponds to observed interactions and the upper right triangle represents simulated contacts using attraction-repulsion maps learned from chromosome 2 of Kc167 cells.

As *Drosophila* only has two autosomes of significant size, we simulated each chromosome with the attraction-repulsion maps of the other. The resulting simulations largely reproduce the compartmental patterning of these chromosomes (Figure 5B, C and Supplemental Figure 5A). Moreover, high resolution maps indicate that the simulation reproduces short range interactions within compartmental domains. This indicates that, just as in humans, these chromatin features or others strongly correlated with them drive the formation of compartmental domains and their interactions. These observations suggest a universal model of compartmentalization in which the fundamental underlying forces driving this process are conserved between organisms.

## Discussion

Interphase chromosomes of vertebrates are generally thought to be organized into large domains whose sequences can be in one of two states--active or inactive (Lieberman-Aiden et al., 2009). Within these large domains are smaller TADs (Dixon et al., 2012), some of which are flanked by CTCF sites in convergent orientation, and therefore correspond to CTCF loops formed by cohesin extrusion, whereas others lack CTCF at their boundaries and are thus formed by different, unknown mechanisms (Rao et al., 2014). Smaller domains termed sub-TADs can also be observed within TADs (Phillips-Cremins et al., 2013). Results reported here address the question of what the unit of eukaryotic chromosome organization is and how we can explain its formation using known biochemical and biophysical forces operating in the nucleus. Answers to this question provided by chromosome conformation studies should account for results obtained using microscopy or biochemical approaches. However, results from Hi-C studies seem to contradict well-established concepts in nuclear biology. First, non-transcribed regions of the genome do not simply interact with each other, as one would conclude from the checkerboard pattern observed in Hi-C heatmaps representing long-range interactions among sequences located in the B compartment. Rather, non-transcribed regions in the genome contain either H3K27me3/PCR1/PCR2, H3K9me3/HP1, or lack either of these histone modifications. Immunofluorescence localization experiments show that Pc-containing regions interact with each other to form Pc bodies, both in *Drosophila* and mammals (Bantignies et al., 2011; Lanzuolo et al., 2007; Li et al., 2011; Tolhuis et al., 2011). The same is true for regions containing H3K9me3 and HP1 that form chromocenters (Huang et al., 2010; Spierer et al., 2005) and actively transcribed regions to form transcription factories (Jackson et al., 1993; Mitchell and Fraser, 2008). Furthermore, recent results indicate that multivalent proteins present in these three types of genomic regions are able to form biomolecular condensates by liquid-liquid phase separation (Boehning et al., 2018; Larson et al., 2017; Strom et al., 2017; Tatavosian et al., 2019). Therefore, compartmental domains and their interactions detected by Hi-C must be more complex than is generally assumed and this complexity must reflect existing observations from microscopy and biochemistry.

Results presented here reveal an underappreciated diversity of compartmental domains, the conserved forces underlying their establishment, and the long-range interactions responsible for the formation of nuclear compartments. Multiple independent forces organize the genome into compartments. Transcriptional activity and H3K9me3 correlate with the strongest of these forces. H3K27me3, while weaker, also correlates directly with compartmental patterns. Finally, sequences lacking any of these histone modifications and enriched in H3K9me2 represent a fourth class of sequences that form their own independent compartment. The dynamic nature of these chromatin features across cell types allows for dynamic compartmentalization during cell differentiation. While both transcriptional activity and H3K9me3 have been previously reported as highly correlated with the formation of nuclear compartments, here we show that each of the biochemical forces associated with the presence of these histone modifications drives compartmentalization independently and can do so in the absence of the other (Falk et al., 2019; Lieberman-Aiden et al., 2009). Previous in-depth Hi-C analyses of GM12878 cells suggested a unique H3K9me3-correlated compartmentalization of chromosome 19 and categorized it as a subcompartment of inactive B chromatin (Rao et al., 2014). Here we show that this compartment, while unique in GM12878, is widespread in other cell lines and is the strongest compartmentalizing force wherever large domains of H3K9me3 are found. On chromosomes where large H3K9me3-rich domains exist, the segregation between these domains and the rest of the H3K9me3-poor chromosome represent the strongest feature of the Hi-C maps. As such, using PCA to delineate binary compartments on the Pearson correlation maps divides the genome into H3K9me3-rich and poor, rather than along the expected lines of transcriptional activity. The inclusion of H3K9me3-poor transcriptionally inactive regions into the A compartment defined by a binary segregation of PC1 in these cells leads to incompatible and confusing definitions of genomic compartments between cell lines and tissues.

While PCA is a powerful tool to investigate compartments, the PC1-based canonical binary classification of compartments is inadequate to represent the true compartmentalization of the genome. We suggest shifting away from naive unsupervised classification techniques in single cell lines to a categorization informed by the breadth of organizational diversity seen across human samples, which will improve the accuracy and generalizability of future studies. Our findings fit a model of compartmentalization where attraction of similar chromatin states drives interactions between them to the exclusion of other chromatin types. The dependence of H3K9me3 compartmentalization in GM12878 cells on the levels of this modification, where small discrete peaks fail to strongly compartmentalize through most of the genome while large domains seen on chromosome 19 do, suggests the existence of a compartmentalization threshold. The forces driving compartmentalization must overcome the entropy of mixing in the nucleus and, therefore, some minimum quantity of compartmentalizing signal must exist below which the attractive forces at work are insufficient to overcome entropy. We propose that the distinctive patterns of H3K9me3 in GM12878 cells reflect this threshold where the discrete ChIP-seq peaks seen in most of the genome fail to strongly compartmentalize while in the same nuclear environment the larger domains present in chromosome 19 do. As further evidence of the quantity-dependent nature of compartmentalization, several chromosomes in HCT-116 cells possess both weak and strong H3K9me3 domains, which correspondingly compartmentalize weakly and strongly. These findings suggest that compartmentalization is more accurately represented as a continuum, where each sequence is driven to interact according to the strength, corresponding to the quantity, of chromatin forces driving it, rather than discrete chromatin types.

Our model provides insight into the fundamental mechanisms of genome 3D organization. That the compartmentalization of the genome can be predicted using just the distribution of a few histone modifications strongly implies that these chromatin features are either directly or indirectly responsible for the spatial separation of chromatin types in the genome. Our finding that the patterns of attraction and repulsion are largely consistent between cell types with divergent compartments shows that these underlying forces behave consistently and that the fundamental forces shaping chromatin organization are steady between cells. The ability of the same model to reproduce the organization of *Drosophila* chromosomes suggests that the attraction and repulsion of chromatin by the independent contributions of compartmentalizing forces may be a universal driver of compartmentalization across Animalia. Taken together, our results reshape our understanding of human compartments from largely static, binary classes to a highly dynamic and quantitative continuum. The widespread use of canonical A/B compartmentalization is largely a product of the pioneering work done in GM12878, but it does not reflect the epigenetic diversity of human cell types. Informing future analyses of compartments with this understanding and approaching compartmental organization from the perspective of chromatin state driven attraction and repulsion will allow for reproducible and comparable definitions of compartments across human tissues and beyond to other organisms.

## STAR★Methods

### Key Resources Table

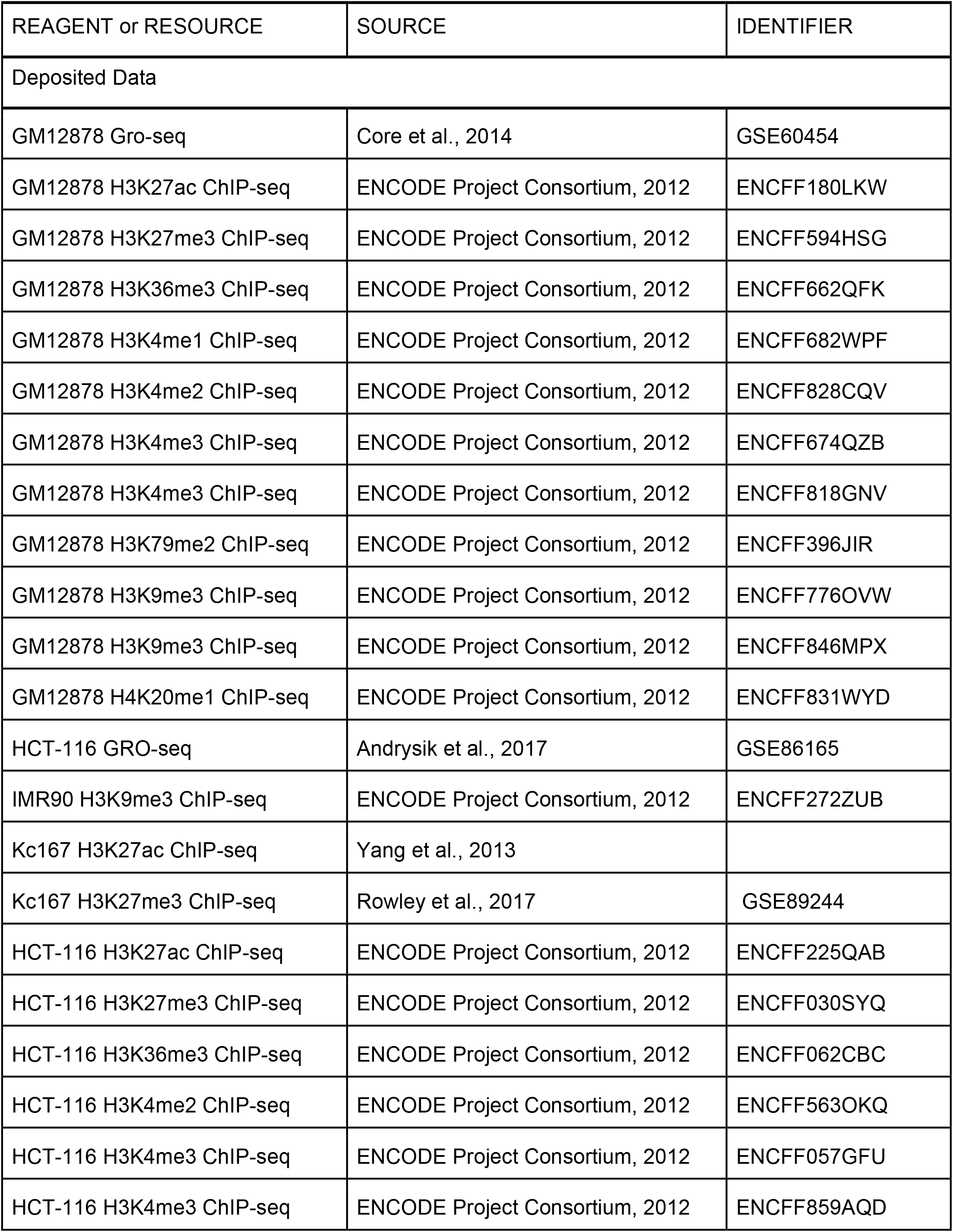
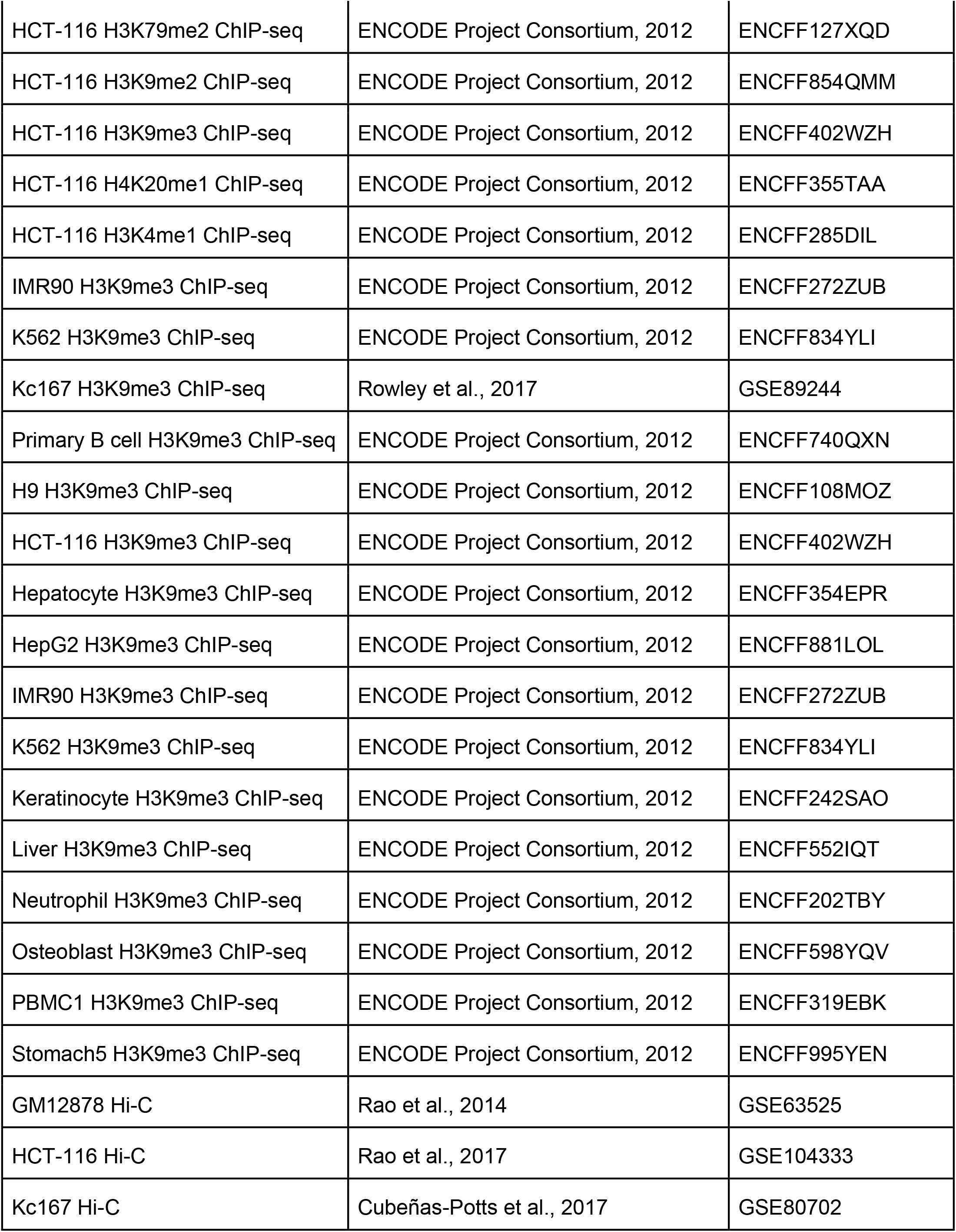
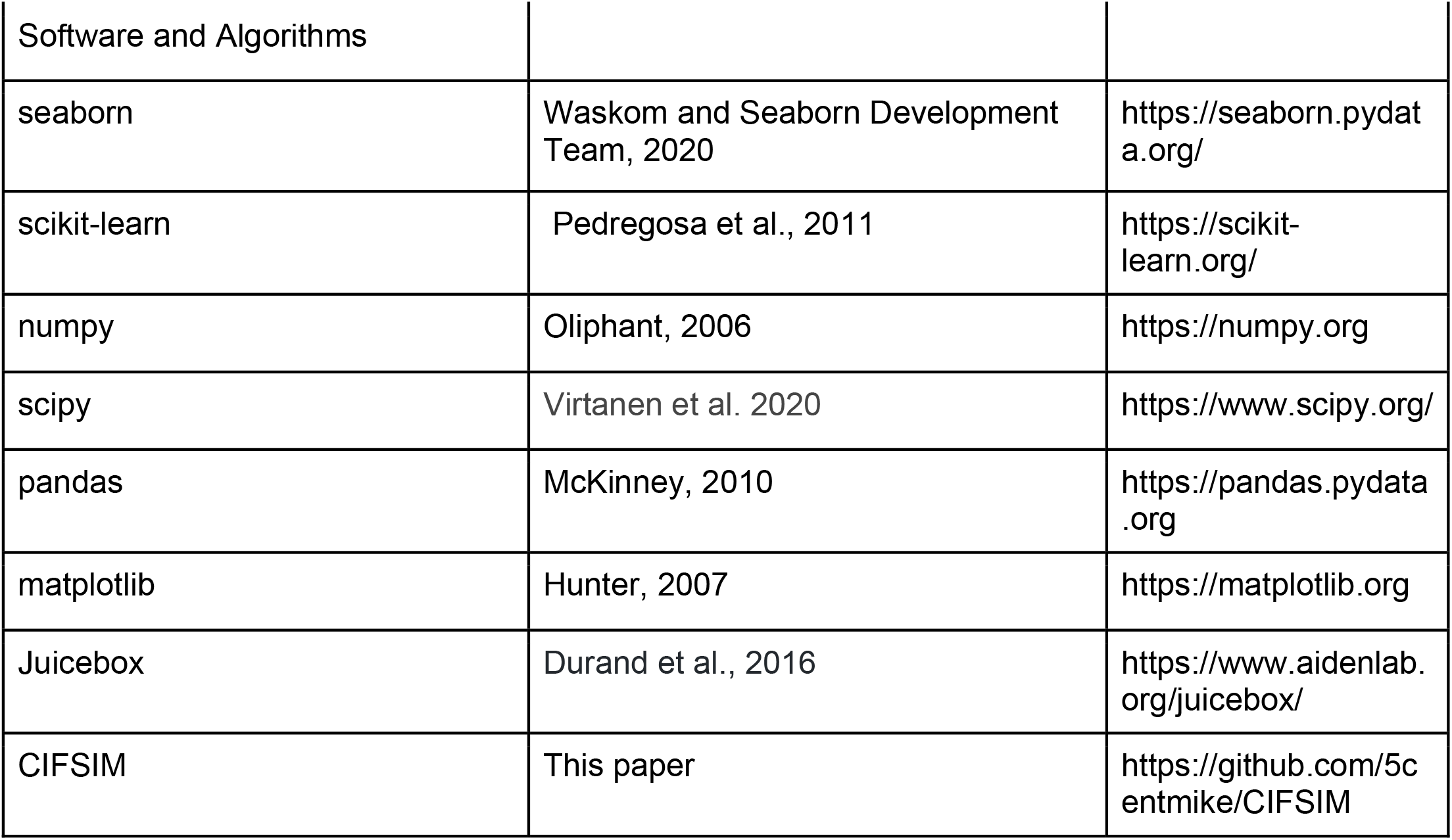

### Resource Availability

This study did not use unique resources.

### Lead Contact

Further information and requests for resources and reagents should be directed to and will be fulfilled by the Lead Contact, Dr. Victor G Corces.

### Materials Availability

This study did not generate new unique reagents.

### Data and Code Availability

The code generated during this study is hosted on GitHub: https://github.com/5centmike/CIFSIM/.

This study did not generate new unique datasets.

## Method Details

### ChIP-seq quantile normalization

Two replicates of ChIP-seq data were combined in bigwig format and the fold-change over control was determined. 100 kb bins with no reads mapped in any ChIP-seq were excluded from analysis. The remaining bins were normalized into twenty discrete quantiles according to their signal compared genome-wide.

### Hi-C Quality Control

Hi-C maps generated from reads with quality score >Q30 were used. Genomic bins were removed from the maps and all subsequent analyses according to several criteria calculated for each chromosome as follows. Bins removed from ChIP-seq quantiles were also removed from the Hi-C. Bins with a total read sum greater than 3 standard deviations above or less than 3 standard deviations below the average bin read sum were dropped. Bins with non-zero interactions with bins with non-zero interactions greater than 3 standard deviations above or less than 3 standard deviations below the average bin were dropped.

### Hi-C Normalization and Pearson correlation

Hi-C maps were balanced using Knight-Ruiz normalization. For some analyses such as Pearson correlation, the Hi-C maps were distance-normalized by dividing each interaction by the average of all interactions at that distance. This produces an observed/expected value for all interaction bins. Pearson correlations of Hi-C maps were then generated from these distance normalized matrices.

### Principal Component Analysis and Compartment calls of Hi-C data

Principal Component Analysis was performed on the chromosome Pearson correlation maps. The first principal component (PC1) is defined as the eigenvector with the largest eigenvalue. All bins with positive values in PC1 were assigned to one compartment while all negative values were assigned to the other. The compartment with the largest enrichment for Gro-seq signal was defined as the A compartment and the other one as B.

### k-means clustering of Hi-C data

To dissect the complex compartmentalization of the genome based on Hi-C information, we employed the unsupervised k-means clustering algorithm to identify clusters in GM12878 and HCT-116 cells. We used MiniBatchKMeans from scikit-learn’s machine learning library with varying numbers for k depending on the features of the given chromosome. Clustering was performed on the Pearson correlation maps of each chromosome separately and clusters were identified as A, B, C, or D by their enrichments for H3K27ac, H3K27me3, H3K9me3, and H3K9me2, respectively.

### Compartmentalization by Independent Forces to Simulate Interaction Maps

To enable comparison across cell types and experiments, we first binned all epigenetic signals into quantiles at 100 kb resolution. For each normalized signal, the model learned, using a Maximum Likelihood Estimation approach, an attraction-repulsion relationship for each pair of quantiles. This attraction-repulsion mapping effectively represents the average enrichment or depletion between all bins with the corresponding level of signal. The model then predicts the number of reads at each bin by summing the attraction-repulsion scores for each signal and multiplying by a constant distance factor to account for the power law decay of genomic interactions.

We model the compartmentalization of the genome as the independent contributions of individual 1D epigenetic signals and use a machine learning method, Maximum Likelihood Estimation, to learn the relationship between the signals and interaction frequencies in the Hi-C map. In this way we hope to quantify the nuclear forces driving compartmentalization. Our model treats each epigenetic signal as an independent but additive effect on Hi-C interaction frequency according to the equation:

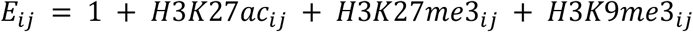

where the expected interaction frequency E between any two genomic loci i and j is the sum of the weight effect of each signal. Using Maximum Likelihood Estimation we find the optimal values of the weights of each signal such that the expected result E is as close to the observed value as possible. This produces a trained model that can reproduce the 3D organization of the genome from 1D epigenetic signals by quantifying the expected contributions of each correlated compartmentalizing force.

We use the Maximum Likelihood Estimation approach to optimize the values of a vector **β** where each entry in **β** corresponds to an entry in the attraction-repulsion maps such that for all possible pairs of quantiles for each of the three chromatin signals H3K27ac, H3K27me3, and H3K9me3, there is a corresponding weight in **β**. We then construct a sparse matrix X where each row corresponds to an interaction bin in the flattened Hi-C matrix and each column a weight in **β**. Each row in X is zeros except in the 3 columns corresponding to the entries in **β** that describe the pair of signal quantiles of genomic bins. If we take an observed Hi-C map which has been normalized for distance by dividing by the expected value at each distance, which we term y, the approximation of the model of the normalized observed map is then:

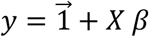

Treating the normalized interaction frequency as a normal distribution we derive a likelihood function the log of which we will maximize to optimize the weights of **β** in a Maximum Likelihood Estimation approach:

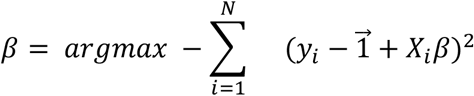

where N is the total number of unique interacting bins in the linearized Hi-C matrix excluding interactions of each genomic bin with itself. To optimize the parameters of **β** we iteratively solve starting with initial values of 0 for all weights in **β**. We use the Newton-Raphson method to update the weights by gradient descent. With each iteration we account for one final feature of the Hi-C matrix. Due to the colocalization of compartmentalizing features along the chromosome, the frequency of intra-compartmental interactions is enriched at short ranges. Two genomic bins within one Mb are more likely to be enriched for compartmental interactions than two bins many Mb apart. This bias leads to aberrant distance normalization and, thus the distance normalization is updated after each iteration to account for the average compartmental enrichment the model predicts at each distance. The distance normalization is divided by the average enrichment so that it more accurately reflects the true effect of distance on interaction frequency.

### Simulation of *Drosophila* Hi-C data

The simulation of *Drosophila* Hi-C data works identically to the simulation described above for human data except for a limit on the maximum size of bins that are considered for the analysis. We excluded all interaction bins further than 2 Mb apart as the compartmental signal beyond this distance was substantially weaker.

### Pearson Correlation Analysis of simulated data

Simulations were generated at 100 kb resolution using the average of the attraction-repulsion maps learned from every chromosome except the one being simulated. We quantified the accuracy of the model using the Pearson correlations between the observed and simulated maps after dividing by the average distance. Due to the power law decay of interaction frequency with respect to distance in Hi-C maps, any simulation that accurately reproduces this decay will have an extremely high correlation. As this would not represent the capacity of the model to reproduce compartmental organization we normalized for distance to eliminate the natural correlation driven by the accurate representation of the distance decay.

## Author Contributions

MHN and VGC designed the project and wrote the manuscript. MHN performed all data analyses and designed and implemented the machine learning algorithms.

## Acknowledgements

We would like to thank Anthony Christodoulou and Andrea Trevino for their insights into machine learning. This work was supported by U.S. Public Health Service Award R01 GM035463 from the National Institutes of Health. MHN was supported by NIH T32 GM008490. The content is solely the responsibility of the authors and does not necessarily represent the official views of the National Institutes of Health.

## Supplemental Figures

**Supplementary Figure 1.**
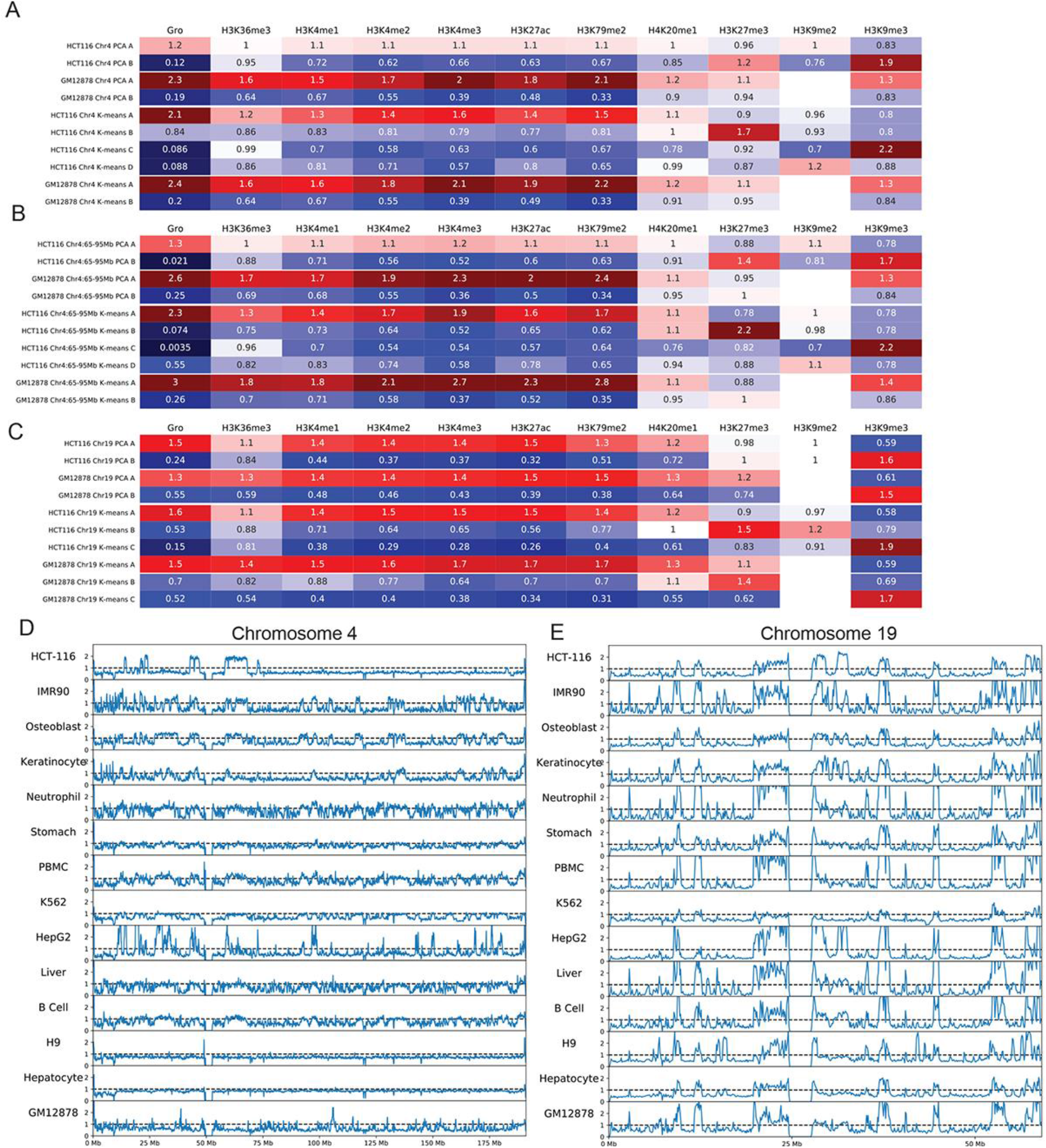
A) Fold-enrichment of each chromatin feature within each compartment defined by PCA or k-means clustering on chromosome 4 of GM12878 and HCT-116 cells. B) Fold-enrichment of each chromatin feature within each compartment defined by PCA or k-means clustering on region 65-95 Mb of chromosome 4 from GM12878 and HCT-116 cells. C) Fold-enrichment of each chromatin feature within each compartment defined by PCA or k-means clustering on chromosome 19 of GM12878 and HCT-116 cells. D) H3K9me3 fold-change over control tracks for a variety of human cell lines and tissues on chromosome 4. E) H3K9me3 fold-change over control tracks for a variety of human cell lines and tissues on chromosome 19.

**Supplementary Figure 2.**
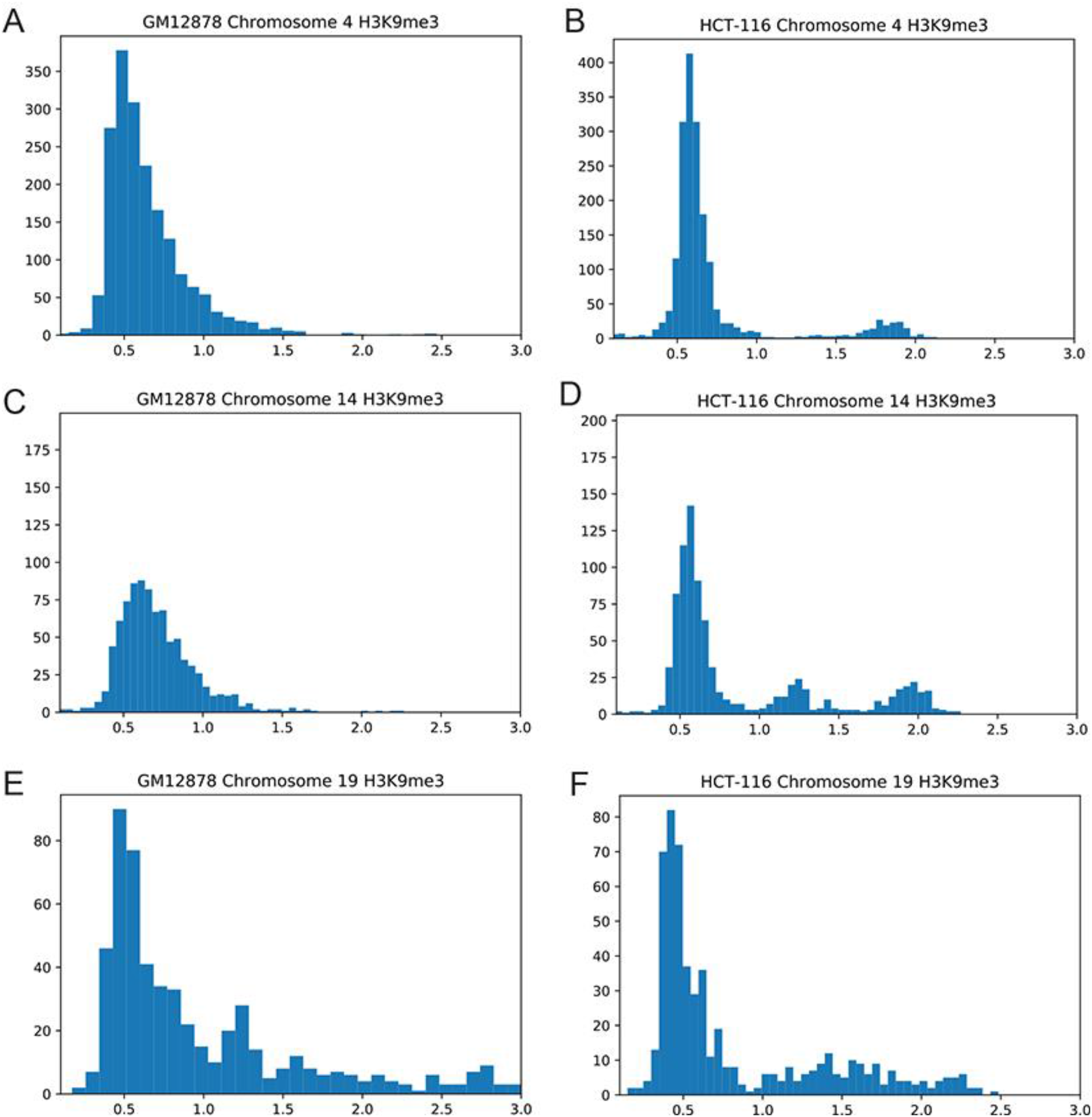
A) 100 kb bin histogram of the distribution of H3K9me3 on chromosome 4 in GM12878 cells. B) 100 kb bin histogram of the distribution of H3K9me3 on chromosome 4 in HCT-116 cells. C) 100 kb bin histogram of the distribution of H3K9me3 on chromosome 14 in GM12878 cells. D) 100 kb bin histogram of the distribution of H3K9me3 on chromosome 14 in HCT-116 cells. E) 100 kb bin histogram of the distribution of H3K9me3 on chromosome 9 in GM12878 cells. F) 100 kb bin histogram of the distribution of H3K9me3 on chromosome 9 in HCT-116 cells.

**Supplementary Figure 3.**
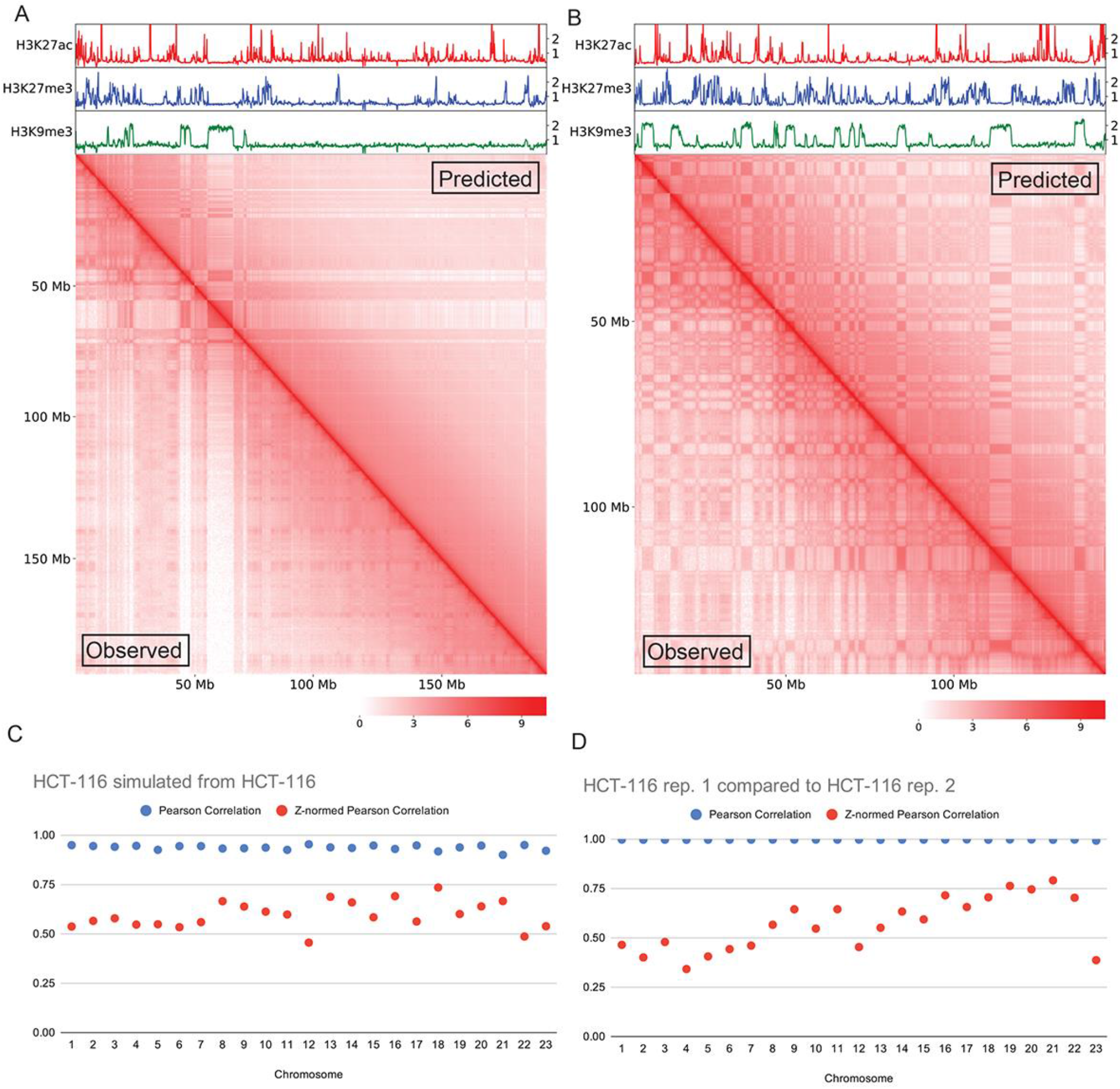
A) Comparison of HCT-116 chromosome 4 logged Hi-C interaction maps. The bottom left triangle represents observed interactions and the upper right triangle represents simulated interactions. B) Comparison of HCT-116 chromosome 8 logged Hi-C interaction maps. The bottom left triangle corresponds to observed interactions and the upper right triangle represents simulated contacts. C) Pearson correlation values comparing the observed and simulated maps for each chromosome of HCT-116 cells. D) Pearson correlation values comparing the two Hi-C replicates from HCT-116 cells.

**Supplementary Figure 4.**
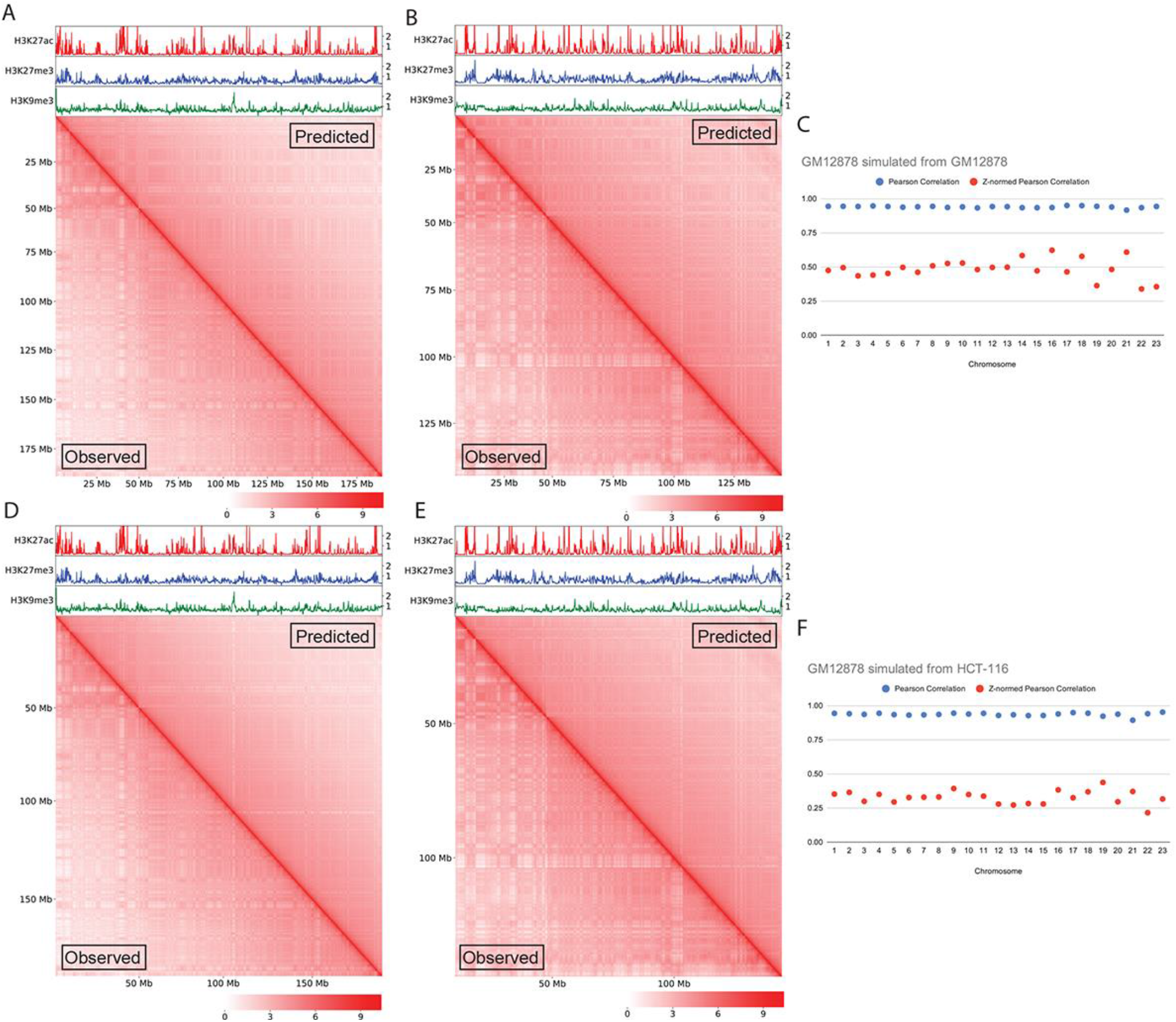
A) Comparison of GM12878 chromosome 4 logged Hi-C interaction maps. The bottom left triangle represents observed interactions and the upper right triangle represents simulated interactions using attraction-repulsion maps learned from GM12878. B) Comparison of GM12878 chromosome 8 logged Hi-C interaction maps. The bottom left triangle represents observed interactions and the upper right triangle represents simulated interactions using attraction-repulsion maps learned from GM12878. C) Pearson correlation values comparing the observed and simulated maps of GM12878 using attraction-repulsion maps learned from GM12878 for each chromosome. D) Comparison of GM12878 chromosome 4 logged Hi-C interaction maps. The bottom left triangle represents observed interactions and the upper right triangle represents simulated interactions using attraction-repulsion maps learned from HCT-116. E) Comparison of GM12878 chromosome 8 logged Hi-C interaction maps. The bottom left triangle represents observed interactions and the upper right triangle represents simulated interactions using attraction-repulsion maps learned from HCT-116. F) Pearson correlation values comparing the observed and simulated maps of GM12878 using attraction-repulsion maps learned from HCT-116 for each chromosome.

**Supplementary Figure 5.**
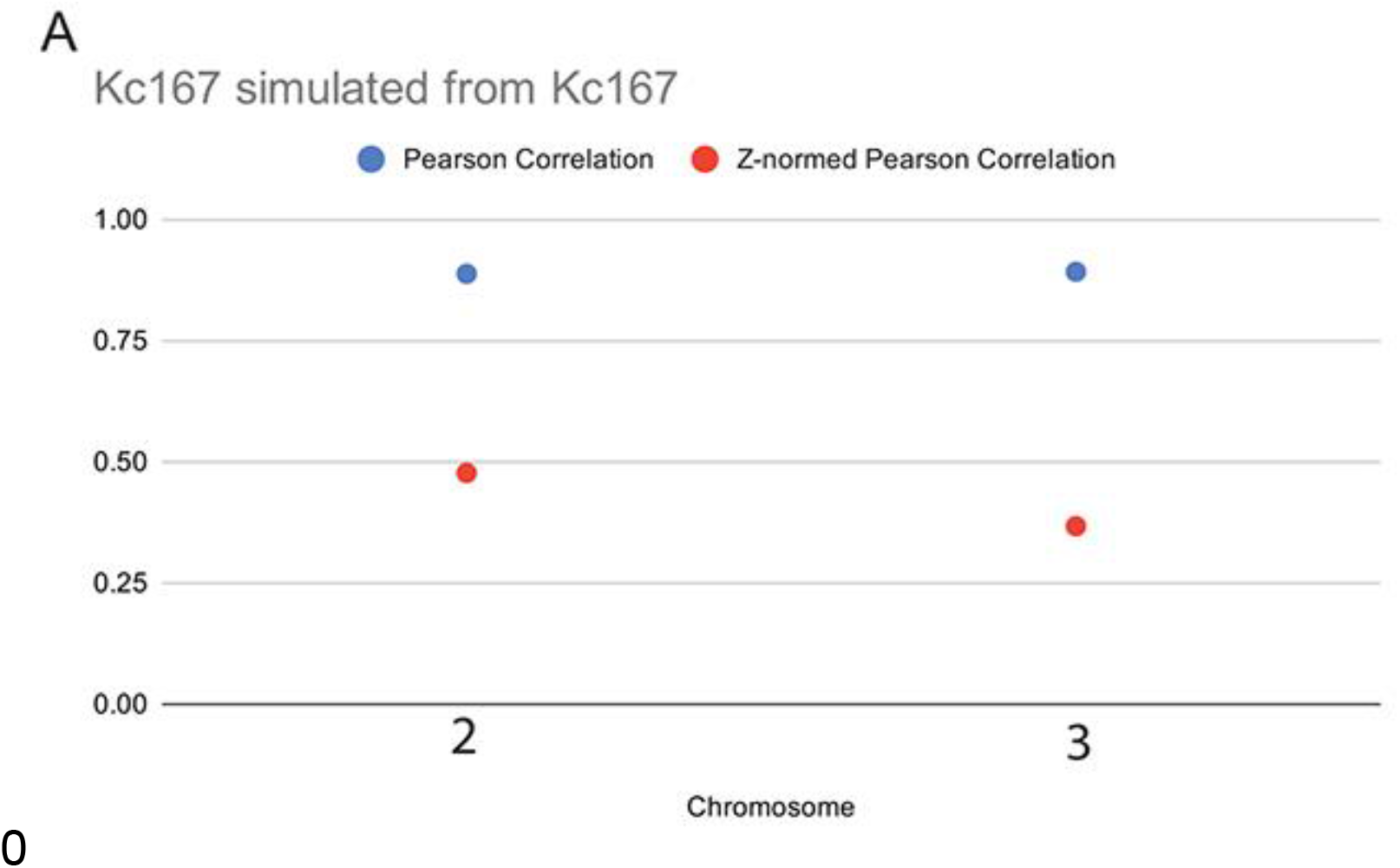
A) Pearson correlation values comparing the observed and simulated maps for each chromosome of *Drosophila* Kc cells.

## References

Andrysik, Z., Galbraith, M.D., Guarnieri, A.L., Zaccara, S., Sullivan, K.D., Pandey, A., MacBeth, M., Inga, A., and Espinosa, J.M. (2017). Identification of a core TP53 transcriptional program with highly distributed tumor suppressive activity. Genome Res.

Annunziatella, C., Chiariello, A.M., Esposito, A., Bianco, S., Fiorillo, L., and Nicodemi, M. (2018). Molecular Dynamics simulations of the Strings and Binders Switch model of chromatin. Methods 142, 81–88.

Banani, S.F., Lee, H.O., Hyman, A.A., and Rosen, M.K. (2017). Biomolecular condensates: organizers of cellular biochemistry. Nature Reviews Molecular Cell Biology 18, 285–298.

Bantignies, F., Roure, V., Comet, I., Leblanc, B., Schuettengruber, B., Bonnet, J., Tixier, V., Mas, A., and Cavalli, G. (2011). Polycomb-dependent regulatory contacts between distant Hox loci in Drosophila. Cell 144, 214–226.

Boehning, M., Dugast-Darzacq, C., Rankovic, M., Hansen, A.S., Yu, T., Marie-Nelly, H., McSwiggen, D.T., Kokic, G., Dailey, G.M., Cramer, P., et al. (2018). RNA polymerase II clustering through carboxy-terminal domain phase separation. Nat Struct Mol Biol 25, 833–840.

Boijja, A., Klein, I.A., Sabari, B.R., Dall’Agnese, A., Coffey, E.L., Zamudio, A.V., Li, C.H., Shrinivas, K., Manteiga, J.C., Hannett, N.M., et al. (2018). Transcription factors activate genes through the phase separation capacity of their activation domains. Cell 175, 1842–1855.e16.

Buckle, A., Brackley, C.A., Boyle, S., Marenduzzo, D., and Gilbert, N. (2018). Polymer Simulations of Heteromorphic Chromatin Predict the 3D Folding of Complex Genomic Loci. Mol Cell 72, 786-797.e11.

Chiang, M., Michieletto, D., Brackley, C.A., Rattanavirotkul, N., Mohammed, H., Marenduzzo, D., and Chandra, T. (2019). Polymer Modeling Predicts Chromosome Reorganization in Senescence. Cell Rep 28, 3212-3223.e6.

Cook, P.R., and Marenduzzo, D. (2018). Transcription-driven genome organization: a model for chromosome structure and the regulation of gene expression tested through simulations. Nucleic Acids Res 46, 9895–9906.

Core, L.J., Martins, A.L., Danko, C.G., Waters, C.T., Siepel, A., and Lis, J.T. (2014). Analysis of nascent RNA identifies a unified architecture of initiation regions at mammalian promoters and enhancers. Nat. Genet. 46, 1311–1320.

Cubeñas-Potts, C., Rowley, M.J., Lyu, X., Li, G., Lei, E.P., and Corces, V.G. (2017). Different enhancer classes in Drosophila bind distinct architectural proteins and mediate unique chromatin interactions and 3D architecture. Nucleic Acids Res 45, 1714–1730.

Dixon, J.R., Selvaraj, S., Yue, F., Kim, A., Li, Y., Shen, Y., Hu, M., Liu, J.S., and Ren, B. (2012). Topological domains in mammalian genomes identified by analysis of chromatin interactions. Nature 485, 376–380.

Dixon, J.R., Jung, I., Selvaraj, S., Shen, Y., Antosiewicz-Bourget, J.E., Lee, A.Y., Ye, Z., Kim, A., Rajagopal, N., Xie, W., et al. (2015). Chromatin architecture reorganization during stem cell differentiation. Nature 518, 331–336.

Durand, N.C., Robinson, J.T., Shamim, M.S., Machol, I., Mesirov, J.P., Lander, E.S., and Aiden, E.L. (2016). Juicebox Provides a Visualization System for Hi-C Contact Maps with Unlimited Zoom. Cell Systems 3, 99–101.

ENCODE Project Consortium (2012). An integrated encyclopedia of DNA elements in the human genome. Nature 489, 57–74.

Falk, M., Feodorova, Y., Naumova, N., Imakaev, M., Lajoie, B.R., Leonhardt, H., Joffe, B., Dekker, J., Fudenberg, G., Solovei, I., et al. (2019). Heterochromatin drives compartmentalization of inverted and conventional nuclei. Nature 570, 395–399.

Huang, H., Yu, Z., Zhang, S., Liang, X., Chen, J., Li, C., Ma, J., and Jiao, R. (2010). Drosophila CAF-1 regulates HP1-mediated epigenetic silencing and pericentric heterochromatin stability. J Cell Sci 123, 2853–2861.

Hunter, J.D. (2007). Matplotlib: A 2D Graphics Environment. Computing in Science Engineering 9, 90–95.

Jackson, D.A., Hassan, A.B., Errington, R.J., and Cook, P.R. (1993). Visualization of focal sites of transcription within human nuclei. EMBO J 12, 1059–1065.

Ladouceur, A.-M., Parmar, B.S., Biedzinski, S., Wall, J., Tope, S.G., Cohn, D., Kim, A., Soubry, N., Reyes-Lamothe, R., and Weber, S.C. (2020). Clusters of bacterial RNA polymerase are biomolecular condensates that assemble through liquid–liquid phase separation. PNAS 117, 18540–18549.

Lanzuolo, C., Roure, V., Dekker, J., Bantignies, F., and Orlando, V. (2007). Polycomb response elements mediate the formation of chromosome higher-order structures in the bithorax complex. Nat Cell Biol 9, 1167–1174.

Larson, A.G., Elnatan, D., Keenen, M.M., Trnka, M.J., Johnston, J.B., Burlingame, A.L., Agard, D.A., Redding, S., and Narlikar, G.J. (2017). Liquid droplet formation by HP1α suggests a role for phase separation in heterochromatin. Nature 547, 236–240.

Li, H.-B., Müller, M., Bahechar, I.A., Kyrchanova, O., Ohno, K., Georgiev, P., and Pirrotta, V. (2011). Insulators, not Polycomb response elements, are required for long-range interactions between Polycomb targets in Drosophila melanogaster. Mol Cell Biol 31, 616–625.

Lieberman-Aiden, E., Berkum, N.L. van, Williams, L., Imakaev, M., Ragoczy, T., Telling, A., Amit, I., Lajoie, B.R., Sabo, P.J., Dorschner, M.O., et al. (2009). Comprehensive Mapping of Long-Range Interactions Reveals Folding Principles of the Human Genome. Science 326, 289–293.

Matera, A.G., Izaguire-Sierra, M., Praveen, K., and Rajendra, T.K. (2009). Nuclear bodies: random aggregates of sticky proteins or crucibles of macromolecular assembly? Dev Cell 17, 639–647.

McKinney, W. (2010). Data Structures for Statistical Computing in Python. Proceedings of the 9th Python in Science Conference 56–61.

Michael Waskom, and Seaborn Development Team (2020). Seaborn (Zenodo).

Mitchell, J.A., and Fraser, P. (2008). Transcription factories are nuclear subcompartments that remain in the absence of transcription. Genes Dev 22, 20–25.

Nora, E.P., Goloborodko, A., Valton, A.-L., Gibcus, J.H., Uebersohn, A., Abdennur, N., Dekker, J., Mirny, L.A., and Bruneau, B.G. (2017). Targeted Degradation of CTCF Decouples Local Insulation of Chromosome Domains from Genomic Compartmentalization. Cell 169, 930-944.e22.

Nuebler, J., Fudenberg, G., Imakaev, M., Abdennur, N., and Mirny, L.A. (2018). Chromatin organization by an interplay of loop extrusion and compartmental segregation. Proc Natl Acad Sci U S A 115, E6697–E6706.

Oliphant, T.E. (2006). A guide to NumPy (USA: Trelgol Publishing).

Pedregosa, F., Varoquaux, G., Gramfort, A., Michel, V., Thirion, B., Grisel, O., Blondel, M., Prettenhofer, P., Weiss, R., Dubourg, V., et al. (2011). Scikit-learn: Machine Learning in Python. Journal of Machine Learning Research 12, 2825–2830.

Phillips-Cremins, J.E., Sauria, M.E.G., Sanyal, A., Gerasimova, T.I., Lajoie, B.R., Bell, J.S.K., Ong, C.-T., Hookway, T.A., Guo, C., Sun, Y., et al. (2013). Architectural Protein Subclasses Shape 3D Organization of Genomes during Lineage Commitment. Cell 153, 1281–1295.

Pirrotta, V., and Li, H.-B. (2012). A view of nuclear Polycomb bodies. Curr Opin Genet Dev 22, 101–109.

Plys, A.J., Davis, C.P., Kim, J., Rizki, G., Keenen, M.M., Marr, S.K., and Kingston, R.E. (2019). Phase separation of Polycomb-repressive complex 1 is governed by a charged disordered region of CBX2. Genes Dev 33, 799–813.

Qi, Y., and Zhang, B. (2019). Predicting three-dimensional genome organization with chromatin states. PLoS Comput Biol 15, e1007024.

Rao, S.S.P., Huntley, M.H., Durand, N.C., Stamenova, E.K., Bochkov, I.D., Robinson, J.T., Sanborn, A.L., Machol, I., Omer, A.D., Lander, E.S., et al. (2014). A 3D Map of the Human Genome at Kilobase Resolution Reveals Principles of Chromatin Looping. Cell 159, 1665–1680.

Rao, S.S.P., Huang, S.-C., Glenn St Hilaire, B., Engreitz, J.M., Perez, E.M., Kieffer-Kwon, K.-R., Sanborn, A.L., Johnstone, S.E., Bascom, G.D., Bochkov, I.D., et al. (2017). Cohesin Loss Eliminates All Loop Domains. Cell 171, 305–320.e24.

Rowley, M.J., and Corces, V.G. (2018). Organizational principles of 3D genome architecture. Nat. Rev. Genet. 19, 789–800.

Rowley, M.J., Nichols, M.H., Lyu, X., Ando-Kuri, M., Rivera, I.S.M., Hermetz, K., Wang, P., Ruan, Y., and Corces, V.G. (2017). Evolutionarily Conserved Principles Predict 3D Chromatin Organization. Mol. Cell 67, 837-852.e7.

Spierer, A., Seum, C., Delattre, M., and Spierer, P. (2005). Loss of the modifiers of variegation Su(var)3-7 or HP1 impacts male X polytene chromosome morphology and dosage compensation. J Cell Sci 118, 5047–5057.

Strom, A.R., Emelyanov, A.V., Mir, M., Fyodorov, D.V., Darzacq, X., and Karpen, G.H. (2017). Phase separation drives heterochromatin domain formation. Nature 547, 241–245.

Tatavosian, R., Kent, S., Brown, K., Yao, T., Duc, H.N., Huynh, T.N., Zhen, C.Y., Ma, B., Wang, H., and Ren, X. (2019). Nuclear condensates of the Polycomb protein chromobox 2 (CBX2) assemble through phase separation. J Biol Chem 294, 1451–1463.

Tolhuis, B., Blom, M., Kerkhoven, R.M., Pagie, L., Teunissen, H., Nieuwland, M., Simonis, M., Laat, W. de, Lohuizen, M. van, and Steensel, B. van (2011). Interactions among Polycomb Domains Are Guided by Chromosome Architecture. PLOS Genetics 7, e1001343.

Virtanen, P., Gommers, R., Oliphant, T.E., Haberland, M., Reddy, T., Cournapeau, D., Burovski, E., Peterson, P., Weckesser, W., Bright, J., et al. (2020). SciPy 1.0: fundamental algorithms for scientific computing in Python. Nature Methods 17, 261–272.

Wang, L., Gao, Y., Zheng, X., Liu, C., Dong, S., Li, R., Zhang, G., Wei, Y., Qu, H., Li, Y., et al. (2019). Histone Modifications Regulate Chromatin Compartmentalization by Contributing to a Phase Separation Mechanism. Molecular Cell 76, 646-659.e6.

Yang, J., Sung, E., Donlin-Asp, P.G., and Corces, V.G. (2013). A subset of Drosophila Myc sites remain associated with mitotic chromosomes colocalized with insulator proteins. Nat Commun 4, 1464.

